# A Poisson-multinomial model for quantifying gene therapy vector integrity using quadplex digital PCR

**DOI:** 10.64898/2026.04.09.717556

**Authors:** Lauren Tereshko, Matthew Ryals, Cullen Mason, Romi Admanit, Jake Gagnon

**Affiliations:** Analytical Development, Biogen, Cambridge, Massachusetts, United States of America; Cencora Global Consulting Services, Conshohocken, Pennsylvania, United States of America; Quantitative Sciences and Development Operations, Biogen, Cambridge, Massachusetts, United States of America

## Abstract

Quantitative characterization of gene therapy vectors is critical for confirming product safety and efficacy. While digital PCR (dPCR) strategies are widely used to estimate sequence integrity, accurate statistical interpretation of multiplexed assays remains a persistent challenge due to limitations of existing analytical models. Here, we describe a quadplex droplet dPCR approach for quantitative characterization of vector integrity across four targets within a single reaction. We adapt a Poisson-multinomial model to enable quantitation of sequence integrity and genetic linkage across all combinations of the four targeted regions. We evaluate both the capabilities and limitations of this approach and provide practical guidance for multiplexed assay design and data interpretation.

## Introduction

Gene therapies can be broadly categorized by their delivery modality, including viral vectors like adenovirus (Ad), adeno-associated virus (AAV), and lentivirus, and non-viral approaches such as lipid nanoparticles (LNP), and naked nucleic acids. While recombinant AAV vectors (rAAV) were initially favored over other modalities, recent advances in non-viral vector delivery have made other modalities competitive alternatives with lower immunogenicity and safety risks and manufacturing advantages over viral strategies [1–3].

Production and successful commercialization of any gene therapy modality pose significant analytical challenges in assessing product quality, safety, and efficacy. For rAAV vectors, production is known to result in heterogeneous populations of capsids (empty, partial, full), complicating assessment of vector identity, purity, and genome integrity [4–7]. Plasmid design and production process parameters can further impact product quality by affecting genome truncation and sequence composition [8–11]. Additional production-derived host cell DNA and plasmid impurities may also be present in the drug substance [4,12]. While separation methods can successfully enrich populations for full capsids, characterization of encapsidated content requires molecular assays to assess sequence integrity, truncation, and genomic composition [4–7,13]. Similar molecular characterization challenges apply to non-viral gene therapies where accurate assessment of purity, identity, and sequence integrity remain critical attributes of product quality [14].

Digital PCR (dPCR) is a valuable tool for gene therapy vector characterization that requires low sample input while being robust, sensitive, and specific. In principle, this technology differs from other types of PCR by partitioning DNA templates into thousands of independent reactions. Samples are diluted to an appropriate range where the majority of partitions will contain one or zero template molecules. Fluorescent primer/probes in each partition produce end-point reactions indicating the binary presence or absence of the targeted sequence [15–17]. Reaction complexity can be increased with instruments capable of detecting multiple fluorescent channels, enabling multiplexed dPCR assays.

Duplex dPCR reactions have been leveraged to identify genetic “linkage” of two regions of interest in nucleic acid templates by targeting the 5’ and 3’ ends of a sequence [18–21]. In principle, this approach can be multiplexed to the nth degree by targeting *n* number of sequences along a template using *n* primer/probe sets, enabling a higher resolution of vector characterization. In practice, the capacity to multiplex assays is limited by the compatibility of fluorophores and the ability of dPCR technologies to resolve fluorescent signals. Presently, the order of multiplexing capability ranges from 6-8 fluorescent channels depending on instrumentation and application [22]. Beyond instrumentation, accurate interpretation of multiplex dPCR data presents a statistical challenge as target number increases. Standard Poisson calculations implemented in dPCR software are primarily intended for absolute quantification of independent targets. While some platforms report linkage estimates, these do not explicitly model the full co-occupancy of multiple linked targets as target number increases. We have previously described use of a Poisson-multinomial framework that provides more accurate linkage estimation in duplexed droplet dPCR reaction data [23]. Here we explore an extension of this model to quadplex reactions to enable quantitation of genetic linkage of DNA sequences between all theoretically possible combinations of four targets.

## Results

### I. Sample preparation and quadplex primer/probe optimization

To determine the accuracy of the adapted quadplex model for estimating DNA template integrity, we linearized lentiviral production plasmid (pLV-CMV-EGFP) templates outside of the viral genome sequence with MfeI to create theoretically 100% “intact” templates.Several candidate primer/probe sets were designed for four targets along the sequence (CMV promoter, EGFP cargo, puromycin resistance, and ampicillin resistance), each with a distinct fluorophore (HEX, Cy5, FAM and Cy5.5 respectively) (Fig 1).

**Fig 1.**
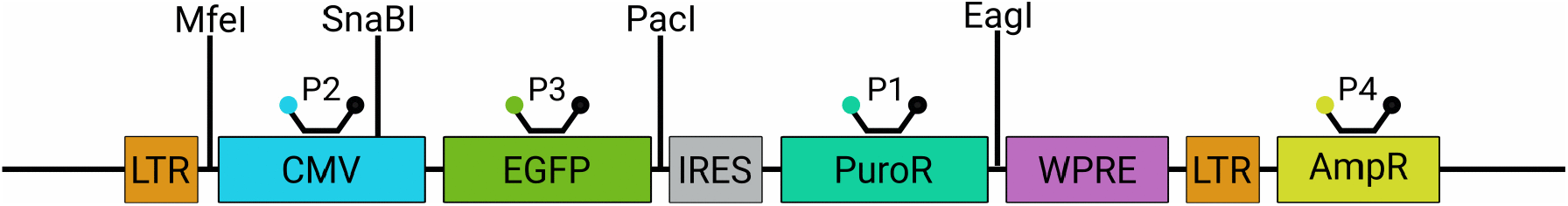
Diagram of linearized pLV-CMV-EGFP with RE sites, target sequences, and primer/probe sets. “PX”= primer probe sets denoted 1:4 according to fluorescent channel order (P1 = FAM, P2 = HEX, P3 = Cy5, P4 = Cy5.5)

Candidate primer probe sets were screened in quadplex droplet dPCR reactions using the Bio-Rad QX ONE instrument. Linearized plasmid samples were serially diluted across a portion of the theoretical quantifiable range of droplet dPCR (0.25-5000 copies/μL), and reactions were assessed for measurement accuracy and minimal crosstalk between fluorescence channels at each dilution [24]. In addition to evaluating probe specificity, these experiments were used to empirically define the concentration range supporting reliable quantitation in quadplex reactions. Software-reported concentrations (copies/µl) were taken from the instrument’s raw data and compared to the expected concentrations derived from the dilution series per channel. While the accuracy of observed versus expected concentration of each target remains stable between 39-5000 copies/μL, increasing variability was seen at lower concentrations (Fig 2). Over-recovery greater than 20% was observed at concentrations ≤20 copies/μL (Table 1). This trend is consistent with the known limitations of droplet dPCR, where at low copy numbers, background fluorescence and stochastic partitioning of a small number of true positive droplets can lead to inflation of Poisson-based concentration estimates [24]. Fig 2 and Table 1 show data from the optimal set of quadplex primer/probes. One- and two-dimensional droplet plots for optimal and alternative quadplex primer/probe panels can be found in S1-S6 Figs.

**Table 1.** Table of average percent recoveries of each channel across dilutions n=2. Gray is outside of ±20% of expected

| Expected copies/μL | AVG % Rec Channel 1 | AVG % Rec Channel 2 | AVG % Rec Channel 3 | AVG % Rec Channel 4 | RSD (all replicates, all channels) |
| --- | --- | --- | --- | --- | --- |
| 5000 | 102.7 | 104.8 | 104.0 | 105.1 | 2.4 |
| 2500 | 102.0 | 103.6 | 103.5 | 103.7 | 3.4 |
| 1250 | 104.3 | 105.8 | 105.6 | 106.1 | 3.6 |
| 625 | 107.1 | 108.7 | 108.5 | 108.9 | 4.6 |
| 312.5 | 107.1 | 108.9 | 108.8 | 109.1 | 1.7 |
| 156.3 | 111.3 | 114.6 | 113.9 | 113.8 | 1.3 |
| 78.1 | 115.7 | 118.5 | 117.5 | 118.5 | 2.2 |
| 39.1 | 113.6 | 117.2 | 115.4 | 118.0 | 3.1 |
| 19.5 | 119.5 | 125.4 | 121.4 | 125.4 | 7.1 |
| 9.8 | 123.9 | 134.3 | 127.9 | 131.3 | 4.4 |
| 4.9 | 123.3 | 145.6 | 125.3 | 139.2 | 9.1 |

**Fig 2.**
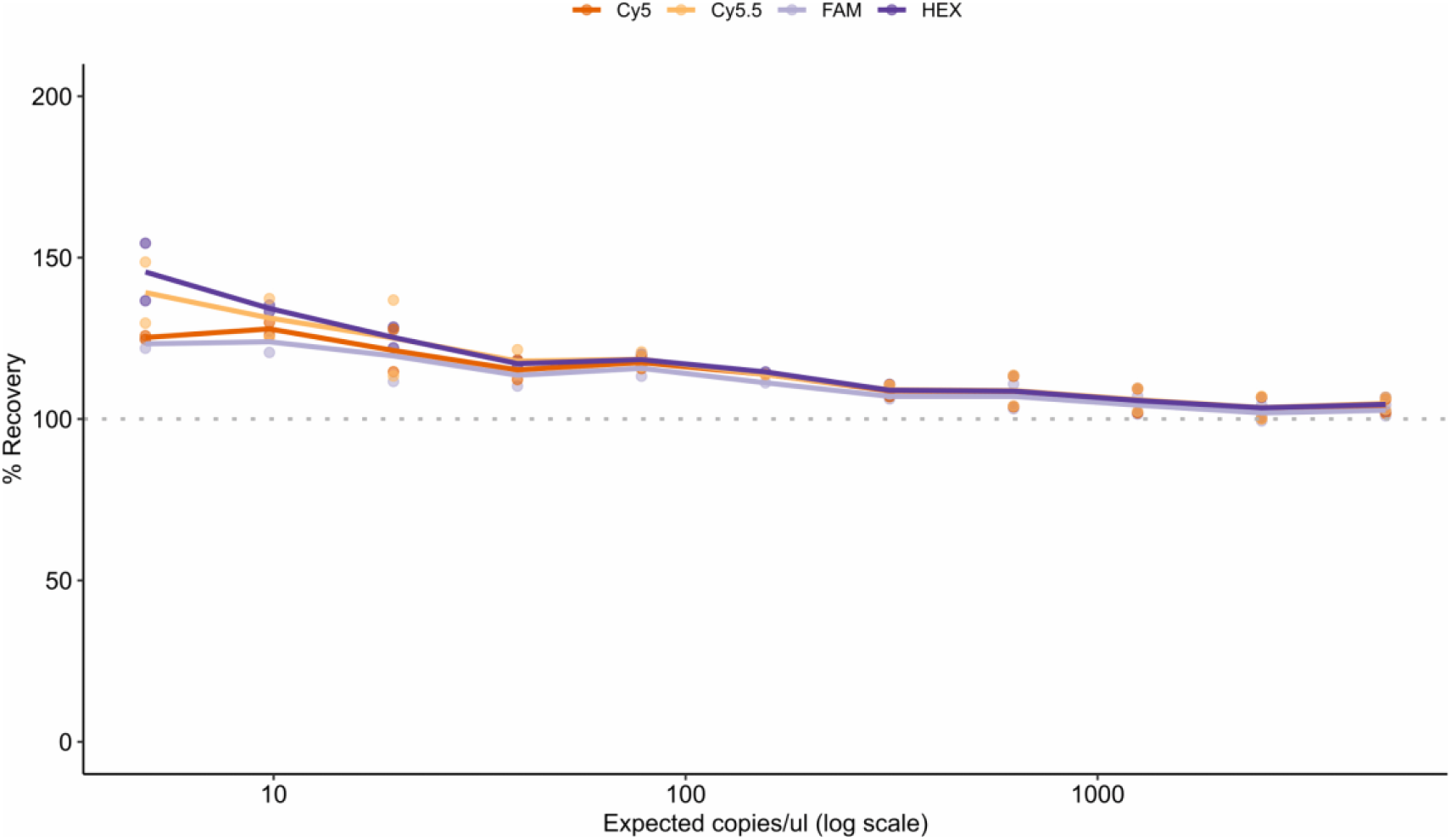
Percent recovery of measured vs expected plasmid concentration per channel n=2. Dashed line denotes 100% recovery. Channels Cy5, Cy5.5, FAM, and HEX correspond to optimized primer–probe sets shown in Fig. 1.

## II. Explanation of quadplex model

Our quadplex model builds upon our previous work in which we described the use of a Poisson-multinomial model to determine the percentage of intact templates in duplex droplet dPCR reactions. The current model begins with a Poisson model where the distribution of the number of templates per droplet is set by the experimentally defined parameter lambda (Formula 1). The likelihood of having *k* templates within a droplet is represented by a Poisson distribution, p(*k*) (Formula 2), where k ranges from 0 to k_max_ and k_max_ is determined by having a cumulative probability ≥ 99%

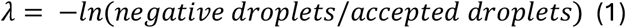

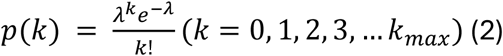

Using the linearized plasmid with four primer/probe targets in Fig 1 as an example and denoting linkage with “_”, there are ten total template types that may be present in a droplet (nine fragments, one intact). Fragmented templates include: CMV, EGFP, PuroR, AmpR, CMV_EGFP, EGFP_PuroR, PuroR_AmpR, CMV_EGFP_PuroR, and EGFP_PuroR_AmpR. Fully intact templates are (CMV_EGFP_PuroR_AmpR). Fixing the number of templates within a droplet to *k*, the four targets and ten linkage combinations can be modeled with a multinomial model with ten species. The model therefore has ten unknown probability parameters. The probabilities of the nine fragmented template species are solved directly, whereas the probability of intact templates is determined by 1 – the sum of fragmented templates (p_CMV_, p_EGFP_, p_PuroR_, p_AmpR_, p_CMV_EGFP_, p_EGFP_PuroR_, p_PuroR_AmpR_, p_CMV_EGFP_PuroR_, and p_EGFP_PuroR_AmpR_).

A multinomial model is defined where each of the *k* templates within a droplet can be one of the ten combinations listed above. The probability of each option is given by: p_CMV_, p_EGFP_, p_PuroR_, p_AmpR,_ p_CMV_EGFP_, p_EGFP_PuroR_, p_PuroR_AmpR_, p_CMV_EGFP_PuroR_, p_EGFP_PuroR_AmpR_, and p_CMV_EGFP_PuroR_AmpR_. For example, p_CMV_ represents the probability that a given template contains the CMV sequence and only the CMV sequence. To derive the ten unknown probabilities, we define *D* as the total number of accepted droplets. We begin by modeling droplets positive for a single target (i.e. CMV amplified by HEX primer/probe). Droplet data from a four-channel instrument results in sixteen unique possible clusters that are either positive or negative for each channel (abbreviated “Ch1-4”). In the Ch1-Ch2+Ch3-Ch4-droplet cluster, we can have combinations such as 1, 2, 3, or more templates positive for the CMV sequence within the droplet. This can be expressed as:

### CMV only model

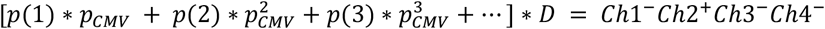 where *p(k)* is the probability of having *k* templates per droplet. This equation can be solved to find the unknown probability quantity, p_CMV_. Similarly, equations can be written for the other single positive droplet clusters to solve for p_EGFP_, p_PuroR_, and p_AmpR_ by considering the EGFP only model, the PuroR only model, and the AmpR only model respectively:

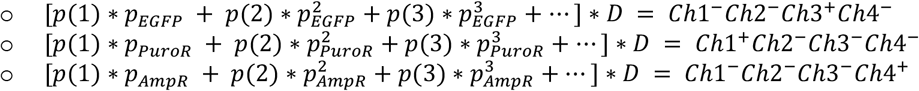

Deriving the probability for the linked targets such as CMV_EGFP involves enumerating the various combinations within the Ch1-Ch2+Ch3+Ch4-droplet cluster:

- 1 template in the droplet: CMV_EGFP

2 templates in the droplet:

- [1 *CMV*, 1 *EGFP*], [2 *CMV*_*EGFP*], [1 *CMV*, 1 *CMV*_*EGFP*], or [1 *EGFP*, 1 *CMV*_*EGFP*]

3 templates in the droplet:

- [3 *CMV*_*EGFP*], [2 *CMV*_*EGFP*, 1 *CMV*],
- [2 *CMV*_*EGFP*, 1 *EGFP*], [1 *CMV*_*EGFP*, 2 *CMV*], [1 *CMV*_*EGFP*, 2 *EGFP*],
- [1 *CMV*_*EGFP*, 1 *CMV*, 1 *EGFP*], [1 *CMV*, 2 *EGFP*], *or* [2 *CMV*, 1 *EGFP*]

The Poisson-multinomial mixture model then yields an expression of the form:

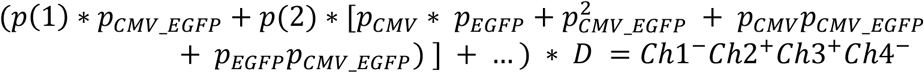

Since p_CMV_ and p_EGFP_ are known from above, p_CMV_EGFP_ can be determined. In a similar manner, expressions can be derived for the remaining unknown variables. However, due to the complexity of these equations, an R script was utilized to enumerate the combinations and to calculate the expressions for p_EGFP_PuroR_, p_PuroR_AmpR_, p_CMV_EGFP_PuroR_, and p_EGFP_PuroR_AmpR_.

## III. Proof of concept with linearized and restriction digested plasmid templates

To assess the quadplex model’s ability to accurately model the proportion of intact templates, eight samples of varying degrees of fragmentation were created by digesting the lentiviral plasmid with combinations of one to four restriction enzymes (REs) (Fig 1, Table 2, S7 and S8 Figs). The theoretically expected fragment types, target linkage, and amplification patterns across the four targets are summarized for each sample in Table 2. Probabilities of amplification per fluorescent channel are denoted as “p” followed by four values (1 or 0) to represent the binary presence (1) or absence (0) of amplification per channel. For instance, the first sample listed Table 2 is linearized outside of the viral genomic sequence by MfeI and is expected to have amplification of all four targets from a single intact linear template (CMV_EGFP_PuroR_AmpR; p1111) as the only theoretically possible combination. In the second sample, MfeI similarly cuts outside the targeted region, while SnaBI bisects the viral genomic sequence into two fragments (CMV; p0100 and EGFP_PuroR_AmpR; p1011).

**Table 2.**
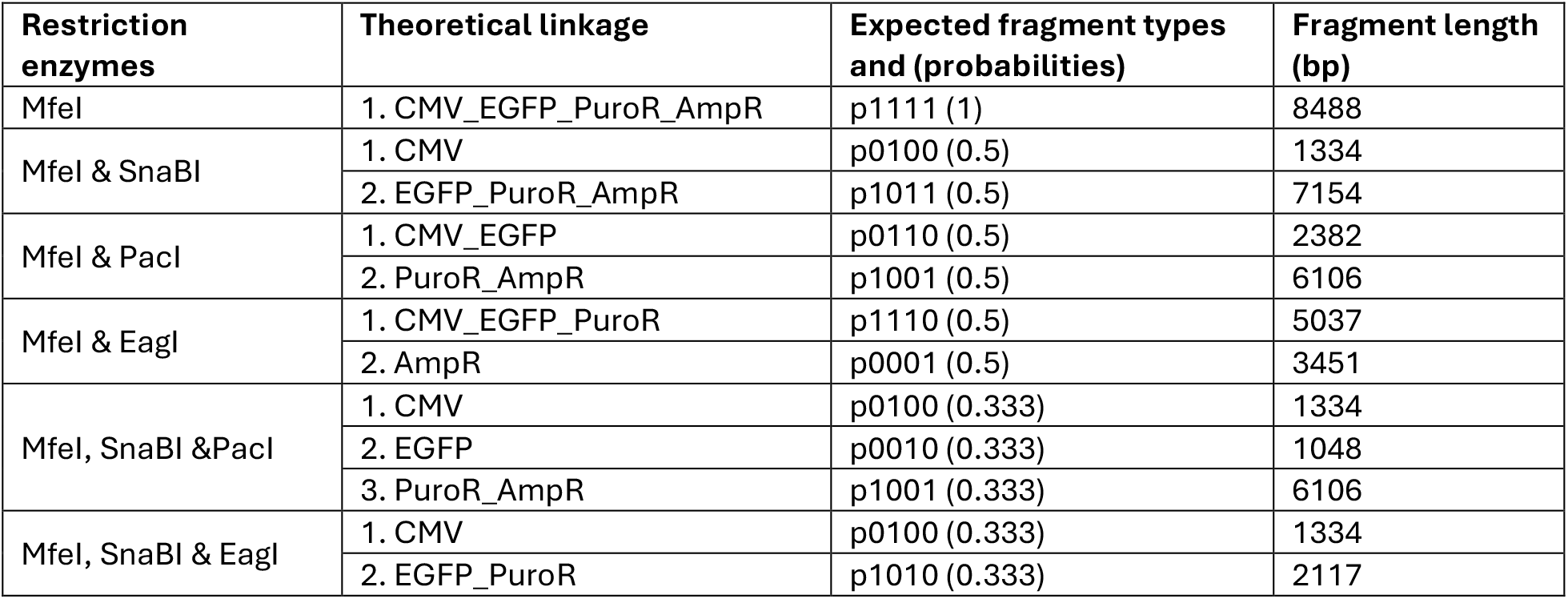
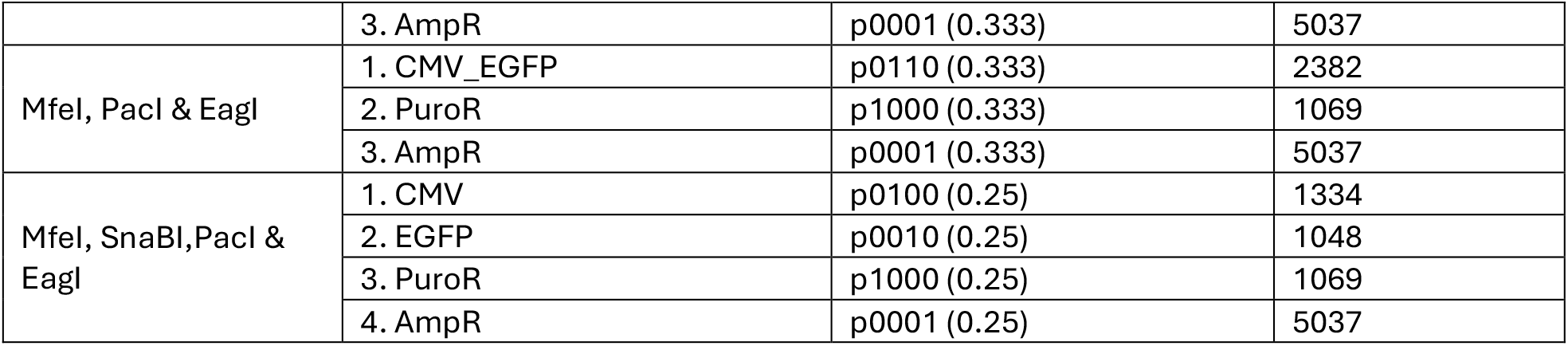
Descriptions of plasmid samples, theoretical linkage and fragment types expected from digestion with REs. Linked targets denoted as being connected by “_”. Binary pattern of expected target amplification per channel is denoted by “pXXXX” followed by the expected probability in parentheses. The XXXX order represents the sequential channel order (with fluorophore and sequence target denoted): Ch1 (FAM, PuroR), Ch2 (HEX, CMV), Ch3 (Cy5, EGFP), Ch 4 (Cy5.5, AmpR)

In the quadplex model, the probability of intact templates (p1111) is not solved for directly and is rather calculated as the remaining probability after estimation of all fragmented template probabilities (Formula 3). Under ideal conditions, the probabilities of all theoretically expected fragmented template species should be between 0 and 1, and sum to 1. In reality, since p1111 is calculated as a derived remainder, negative values can occur if the estimated probabilities of fragmented templates exceeds 1.

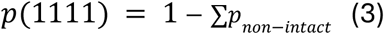

Each sample was serially diluted across 5-5000 copies/μL and run in quadplex reactions. QX ONE data files (Visible Data and Cluster Data exports) were analyzed by the model coded in R, and modeled probabilities were compared to theoretical expectations.

Interestingly, model performance varied with fragmentation complexity, with samples digested into <3 fragments generally exhibiting more accurate probability estimates than samples digested into ≥3 fragments (Fig 3). When plotted as modeled probability vs theoretical concentration and color coded by fragment type, modeled estimates for fully intact linearized plasmid controls (MfeI digests) were consistent with expectations (Fig 3A). To assess the impact of target placement on model estimates, intact controls were also assayed using a second primer/probe panel targeting non-overlapping genomic regions. Comparable intact estimates were obtained using the alternative panel despite reduced assay performance compared to the optimal primer/probe set (S9 Fig).

**Fig 3.**
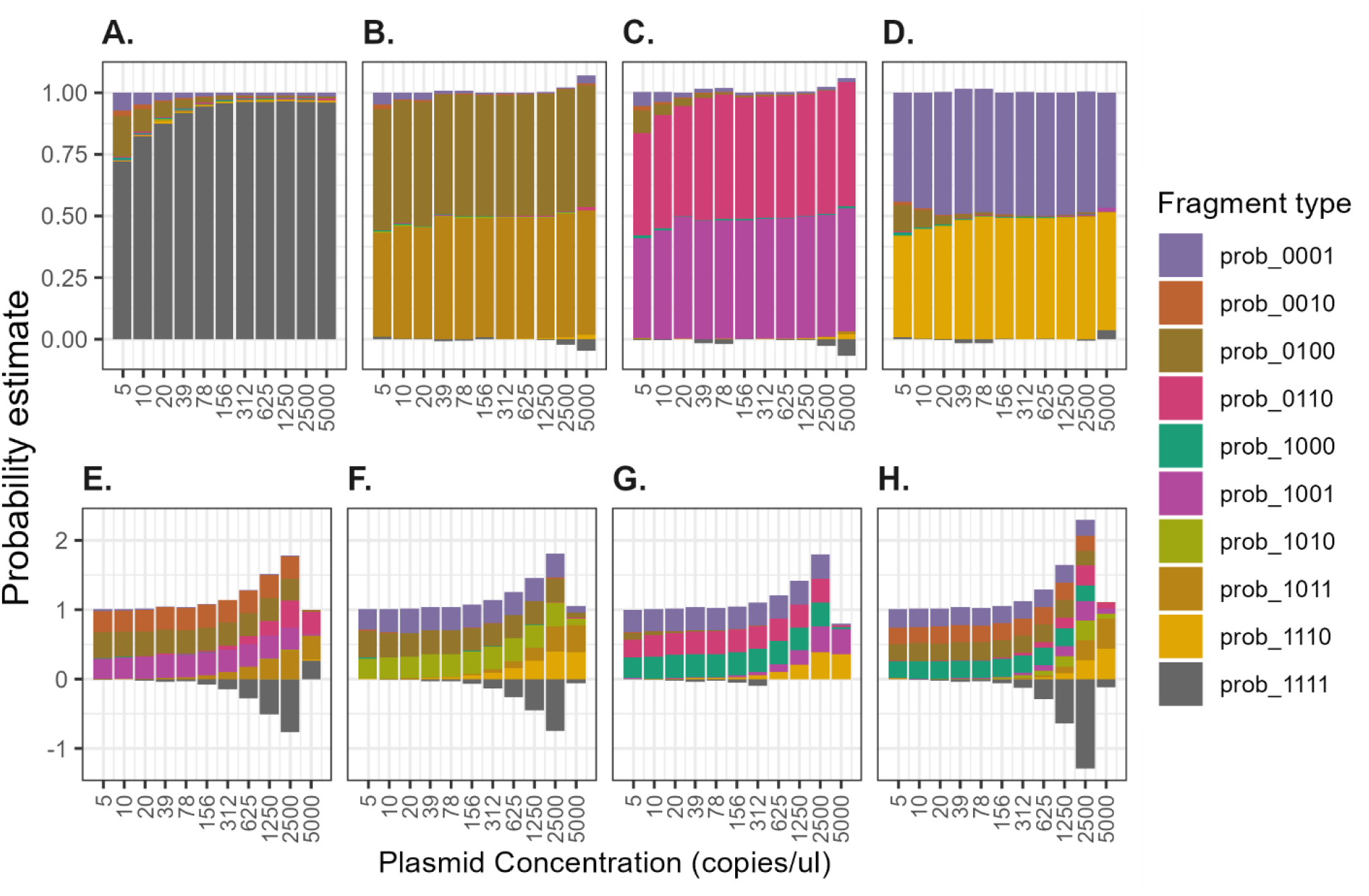
Model results for digested samples described per panel, n=4. A. MfeI, B. MfeI & SnaBI, C. MfeI & PacI, D. MfeI & EagI, E. MfeI, SnaBI & PacI, F. MfeI, SnaBI & EagI, G. MfeI, PacI & EagI, H. MfeI, SnaBI,PacI & EagI

Modeled estimates for samples digested into <3 fragments (by ≤2 REs) were relatively stable across the diluted range, with lower accuracy observed at concentrations below 39 copies/μL (Fig 3A-D). This agrees with the results previously observed in Figure 2. In contrast, modeled estimates for samples digested into ≥3 fragments (by ≥3 REs) increasingly deviated from expected probabilities as template concentrations increased (Fig 3E-H). At concentrations of ≥312 copies/μL, the model also began to over-represent theoretically impossible fragment types for these samples. For example, for concentrations between 5-156 copies/μL, the results for MfeI, SnaBI &PacI (Fig 3E) digested template have the expected proportions (p=0.3333) of the 3 fragment types that are theoretically possible to result from the digestion (p0100, p0010, p1001, as represented by 3 even blocks of color per bar in the bar graph). At concentrations ≥312 copies/μL, however, theoretically unexpected fragment types (such as p0110) begin to have non-zero probability estimates (Fig 3E). Additionally, as these unexpected fragment types become represented, the sum of all probability estimates begins to climb above 1, and as a result, negative estimates for p1111 begin to appear due to over-estimates of theoretically “impossible” fragment types (Fig 3E-H). Furthermore, probability estimates at the upper measured concentration (5000 copies/μL) were inaccurate specifically for templates digested ≥3 fragments (Fig 3E-H, S10 Fig).

## IV. Proof of concept with mixed populations of plasmid digests

Based on the preceding results, we hypothesized that the distributions of fragment size and number may affect the model’s accuracy. At equivalent plasmid-input concentrations,more extensive digestion generates a greater number of independently partitioning template fragments. We therefore hypothesized that samples with higher abundance of smaller fragments could exhibit higher droplet occupancy and reduced model accuracy. To evaluate model performance with heterogeneous template populations, we prepared six sample mixtures containing varying ratios of theoretically intact template (MfeI digests) and bisected template (MfeI & PacI digests) (Table 3). These samples were compared with six mixtures of varying ratios of theoretically intact template and more extensively fragmented template (MfeI, SnaBI, PacI &EagI digests) (Table 4). In this way, the ranges of expected intact probabilities (p1111) overlapped for both mixture sets, while the composition of fragment populations differed. Fig 4A-F shows the integrity estimates of samples with ≤3 expected fragment types are more stable across the dilutional range than samples with ≥4 expected fragments (Fig 4G-L). Probability estimates were least accurate for the samples with the highest proportions of small fragments (Samples 7-9, Fig 4G-I). In these samples, model estimates were more sensitive to sample concentration with substantial deviation from theoretical values at concentrations between 638-5000 copies/μL (Fig 4G-L, S11 Fig).

**Table 3.** Larger fragment sample preparation. Samples prepared as mixtures of MfeI digests (100% intact) and MfeI & PacI (bisected) digests and their expected target amplification probabilities

| Sample Number | MfeI digest (%) | MfeI & PacI digest (%) | Expected fragment type probabilities |  |  |
| --- | --- | --- | --- | --- | --- |
|  |  |  | p1111 | p0110 | p1001 |
| Sample 1 | 0% | 100% | 0 | 0.5 | 0.5 |
| Sample 2 | 40% | 60% | 0.25 | 0.375 | 0.375 |
| Sample 3 | 60% | 40% | 0.429 | 0.286 | 0.286 |
| Sample 4 | 80% | 20% | 0.667 | 0.167 | 0.167 |
| Sample 5 | 90% | 10% | 0.818 | 0.091 | 0.091 |
| Sample 6 | 100% | 0% | 1 | 0 | 0 |

**Table 4.** Smaller fragment sample preparation. Samples prepared as mixtures of MfeI digests (100% intact) and MfeI, SnaBI, PacI & EagI (completely fragmented) digests and their expected target amplification probabilities

| Sample Number | MfeI digest (%) | SnaBI, EagI, PacI & MfeI digest (%) | Expected fragment type probabilities |  |  |  |  |
| --- | --- | --- | --- | --- | --- | --- | --- |
|  |  |  | p1111 | p0100 | p0010 | p1000 | p0001 |
| Sample 7 | 0% | 100% | 0 | 0.25 | 0.25 | 0.25 | 0.25 |
| Sample 8 | 40% | 60% | 0.143 | 0.214 | 0.214 | 0.214 | 0.214 |
| Sample 9 | 60% | 40% | 0.273 | 0.182 | 0.182 | 0.182 | 0.182 |
| Sample 10 | 80% | 20% | 0.5 | 0.125 | 0.125 | 0.125 | 0.125 |
| Sample 11 | 90% | 10% | 0.692 | 0.077 | 0.077 | 0.077 | 0.077 |
| Sample 12 | 100% | 0% | 1 | 0 | 0 | 0 | 0 |

**Fig 4.**
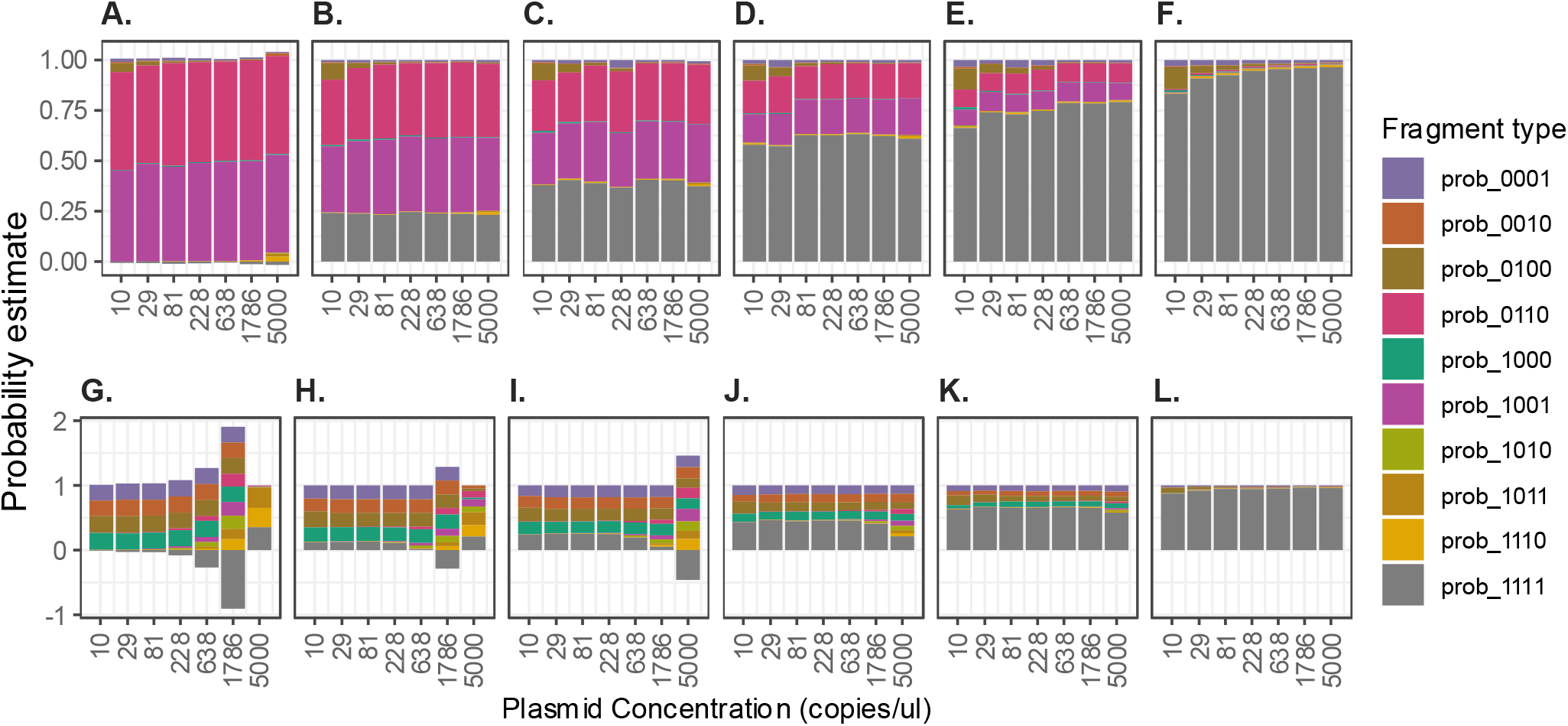
Model results for mixed fragment samples. A-F. results ≤3 fragments (samples 1-6 left to right, top panel), G-L results >3 fragments (samples 7-12 left to right lower panel), n=3.

Our modeling data showed that samples with low or zero empty droplet clusters lead to large lambda (λ) values and inaccurate estimates of intact plasmid. When low or no empty droplets are observed, then the maximum available λ is substituted in modeling that sample, leading to nonideal estimators of fragment probabilities. To evaluate the relationship between occupancy and model performance, intact-template estimates from the fully fragmented samples (MfeI, SnaBI, PacI & EagI digests) in Fig 3H and Fig 4G were transformed and plotted against the fraction of empty droplets ((N-k)/N), the fraction of positive droplets (k/N), and droplet occupancy (λ) (Fig 5A-C). LOESS fitting identified an elbow point estimate at an empty droplet fraction of approximately 27.6% (Fig 5A). These results indicate that, for these plasmid experiments, model performance deteriorated as the fraction of empty droplets decreased and droplet occupancy (λ) increased, highlighting the importance of model performance evaluation with controlled experiments per template type before interpreting model estimates.

**Fig 5.**
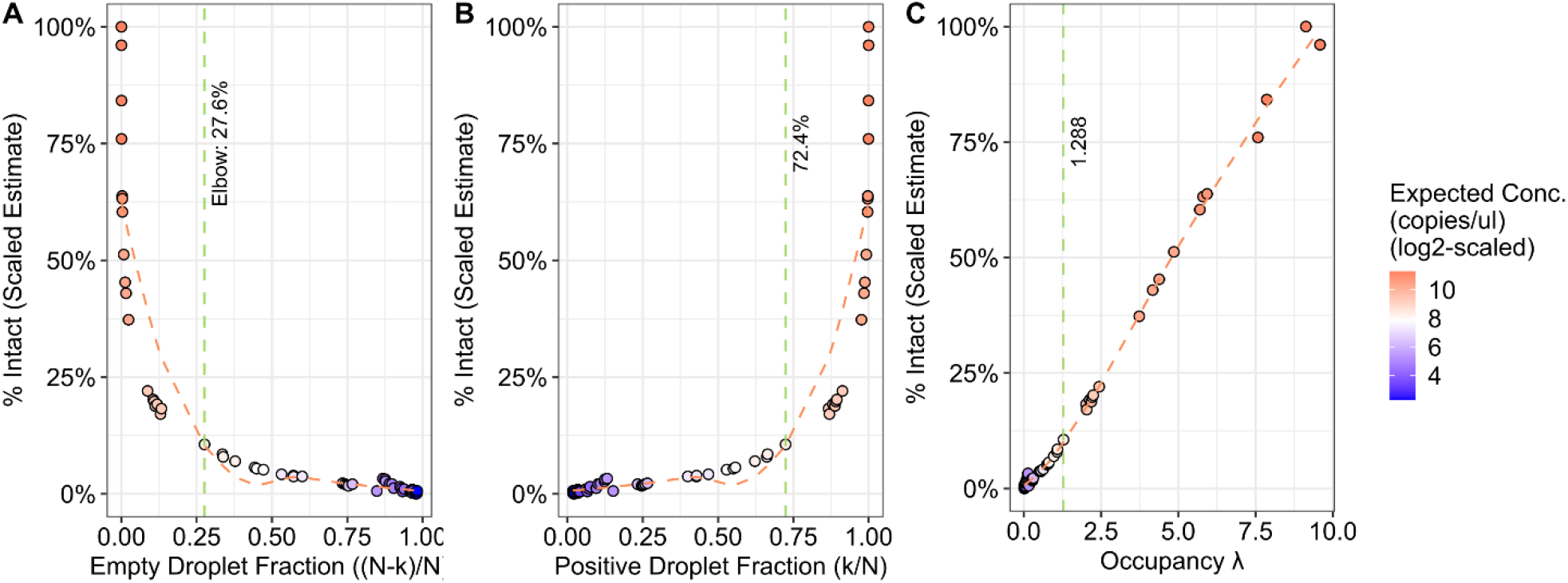
Correlation results of droplet occupancy and model accuracy. A: Scaled model estimates vs empty droplet fraction. B: Scaled model estimates vs positive droplet fraction. C: Scaled model estimates vs λ. Dashed orange lines represent LOESS fits. Green dashed lines represent elbow points.

To further evaluate the model’s performance with heterogeneous template populations, deletion variants of pLV-CMV-EGFP lacking either the CMV promoter (ΔCMV) or WPRE element (ΔWPRE) were generated and mixed with intact parental plasmid at known ratios (S1 Table, S2 Table). Despite <100% recovery of the respective expected p1011 and p1110 fragment types for ΔCMV and ΔWPRE controls, the quadplex model estimates tracked with the expected input compositions across sample mixtures, supporting the ability of the model to distinguish heterogeneous template populations under controlled conditions (S12 Fig, S3 Table).

## V. Applying the quadplex model to extracted rAAV DNA

We next wanted to test the utility of the quadplex model in estimating the integrity of representative viral gene therapy material. As rAAV products are known to be mixed populations of empty, partial, and full capsids, decapsidation of rAAV gene therapy samples results in a heterogeneous mix of fragmented and intact genomic templates.

Decapsidation of rAAV prior to dPCR droplet generation is critical as capsids could possibly contain unlinked targets on separate genome fragments that could result in double- or multi-positive droplets. Decapsidating rAAV templates first ensures that the liberated genomic templates are partitioned independently, allowing true linkage to be assessed.

We first developed and optimized quadplex primer/probe sets for two rAAV gene therapy products (AAV9-BIIB1 and AAV2-BIIB2) using the production plasmid for each product to serve as theoretically 100% intact controls (S13 and S14 Figs). Representative droplet plots for samples and no template controls can be found in S15-S20 Figs. To evaluate the model’s ability to capture differences in the percentage of intact genomic templates, rAAV samples were subjected to degradation by either heat or UV treatment to generate materials spanning a range of fragmentation (Figs 6 and 7 respectively). High heat is known to cause spontaneous hydrolysis of DNA. UV radiation is known to cause DNA damage and UV-C germicidal lamps have been shown to effectively degrade Ad and AAV [25–31].

**Fig 6.**
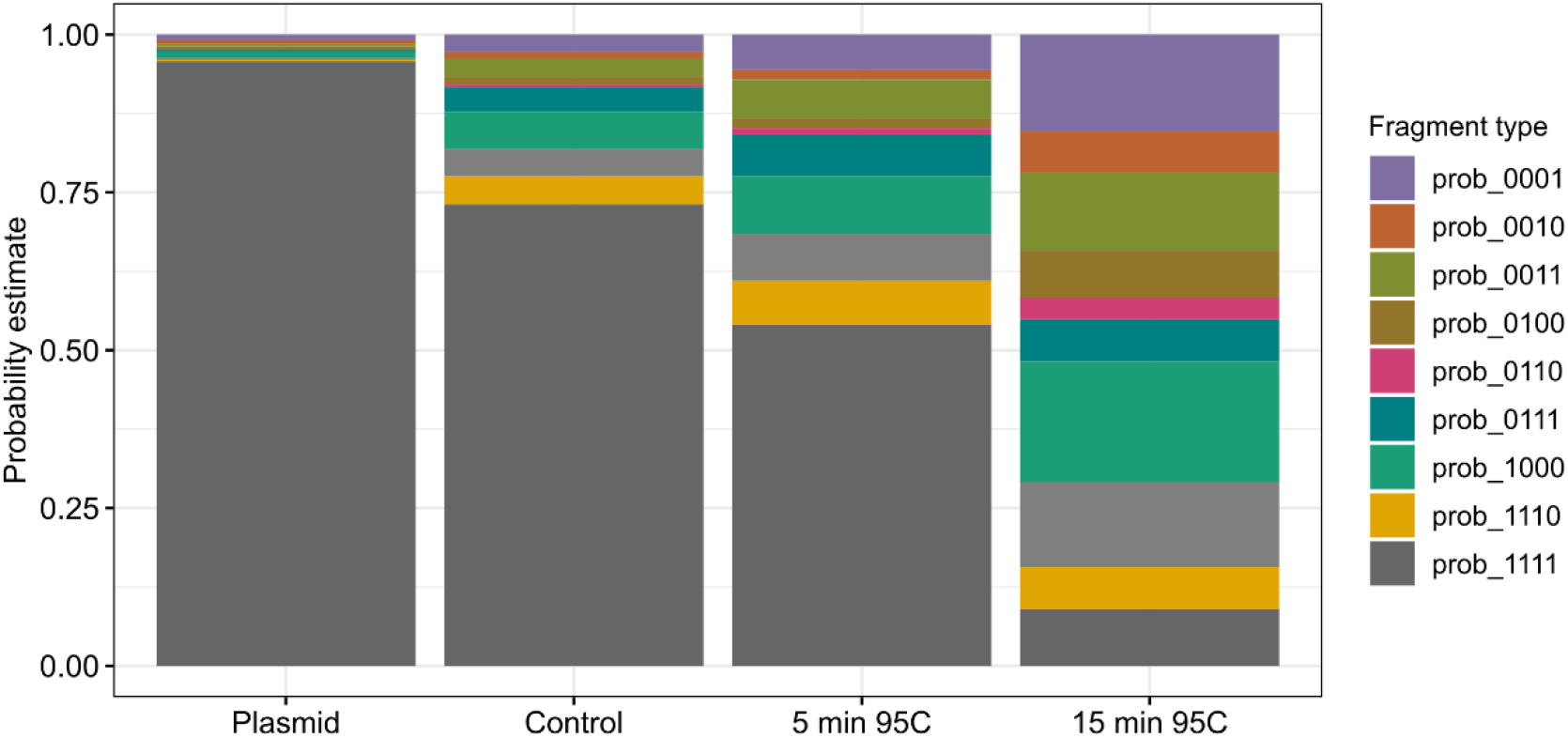
Model captures AAV9-BIIB1 genome degradation from heat treatment (n=4). Average intact probability estimates of plasmid and decapsidated controls and heat samples presented as the grand average of acceptable thresholded concentrations and experimental replicates.

**Fig 7.**
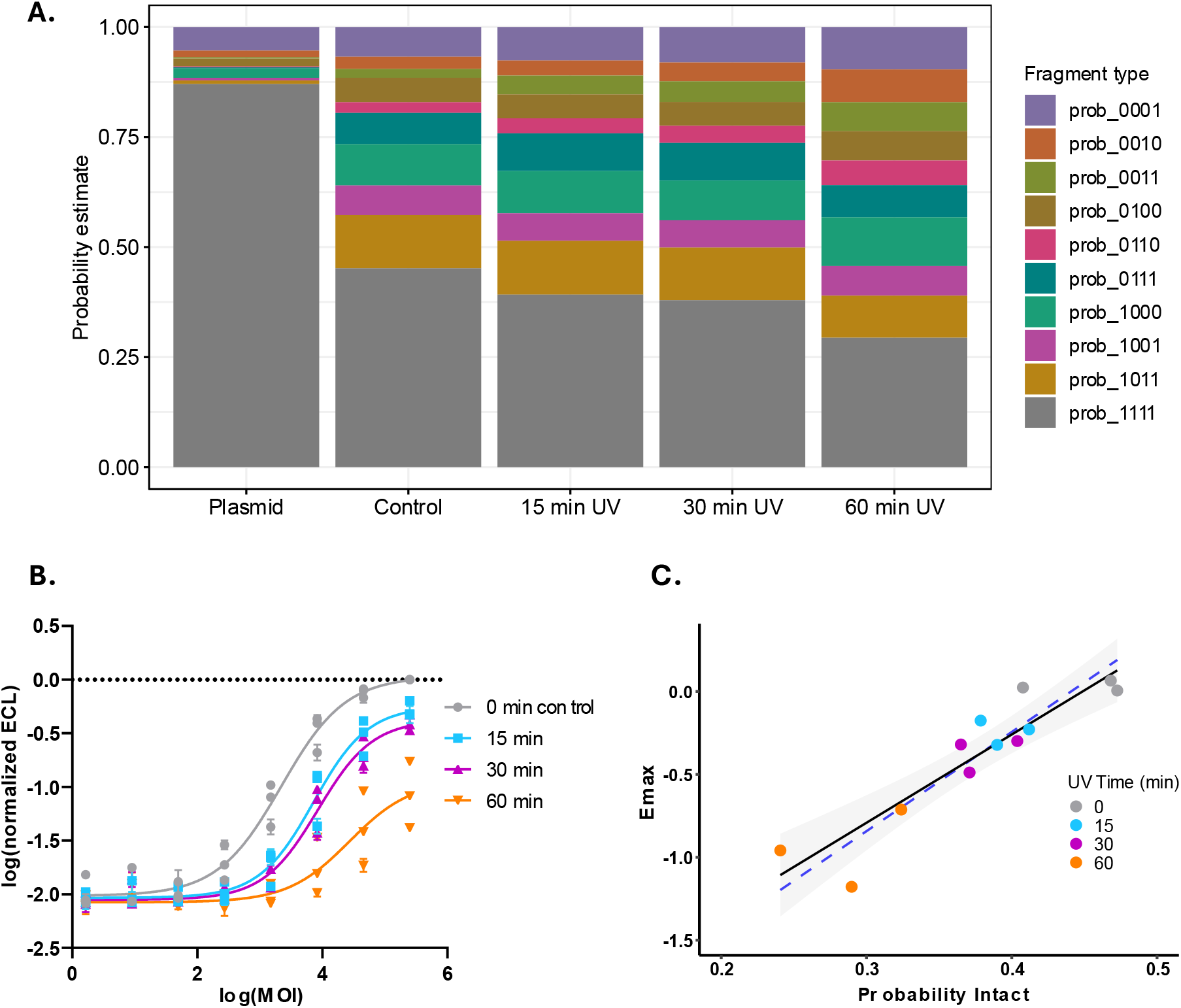
Model captures AAV2-BIIB2 genome degradation from UV treatment (n=3). A: Average intact probability estimates of controls and UV-treated samples presented as the grand average of acceptable thresholded concentrations and experimental replicates. B: AAV2-BIIB2 GOI expression vs multiplicity of infection (MOI), points are color coded by UV treatment duration. C: Correlation of Emax (upper asymptote) with average intact probability estimate of acceptable concentrations per experimental replicate, points are color coded by UV treatment duration. Solid line shows simple linear regression fit with 95% confidence interval shaded in gray, dashed blue line shows linear mixed-effects model fit.

Decapsidated AAV9-BIIB1 material was subjected to either 5 or 15 minutes of 95°C heat. Post-treatment, each sample was serially diluted (5000-20 copies/μL) and run in a quadplex droplet dPCR reaction using the optimized primer/probe sets (S13 Fig). As expected, the average probability estimates for intact templates (p1111) in the plasmid control sample were close to the theoretical value of 1 (0.96); whereas the average estimate for the AAV9-BIIB1 decapsidation control was significantly less than 1 (0.73) due to well-characterized heterogeneity of rAAV capsids. Heat treatment of AAV9-BIIB1 resulted in lower average probability estimates of intact genomes and higher fragment type diversity (Fig 6).

Long-read NGS was performed on a single experimental replicate of control and heat-stressed AAV9-BIIB1 material as an orthogonal approach to quantify genome integrity. As NGS requires purified duplex DNA templates, the single stranded AAV genomes were annealed and purified prior to sequencing. Quadplex droplet dPCR was then performed on the purified duplex samples to enable direct comparison of integrity estimates between methods. Although integrity estimates differed between sequencing and quadplex droplet dPCR, both approaches demonstrated a similar stress-dependent reduction in genome integrity, providing orthogonal support for the degradation trends identified by the quadplex droplet dPCR model (S4 Table and S21 Fig).

AAV2-BIIB2 material was subjected to a time course of UV-C stress (0, 15, 30, or 60 minutes). Post-treatment, samples were serially diluted (5000-20 copies/μL) and quadplex droplet dPCR reactions were run with optimized primer/probes to assess the percentage of intact genomes (S14 Fig). Each sample was also transfected into mammalian cell lines and assayed (in triplicate) for viral cargo protein expression via MSD immunoassay 48 hours later.

As expected, the average probability estimate for intact templates (p1111) in the plasmid control sample was close to the theoretical value of 1 (0.87); whereas the estimate for the AAV2-BIIB2 0-minute UV control was significantly less than 1 (0.45) due to well-characterized heterogeneity of rAAV capsids. UV-C treatment resulted in both lower probability estimates of intact genomes and lower protein expression of the viral gene of interest (Fig 7A and 7B respectively). The reduction in model estimates for intact genomes correlated with reductions in functional output as measured by maximal protein expression (E_max_), supporting that the assay can capture biologically meaningful degradation of gene therapy vectors (Fig 7C, y = 5.32x-2.39, Pearson r = 0.91, p < 0.0001; R^2^ = 0.82). Similar results were observed using a linear mixed-effects model accounting for variation in experimental replicates (Fig 7C, β = 5.98, p < 0.0001).

Variable low-level background noise was observed in no template control wells for this quadplex panel that was higher than what was observed for the pLV-CMV-EGFP and AAV9-BIIB1 panels. Background signal of the WPRE target was most pronounced, suggesting a panel-specific phenomenon. While the observed background did not affect the interpretation of results, it may have contributed to the lower intact-template recovery observed for the corresponding linearized plasmid control (0.87) compared to controls evaluated with other quadplex panels.

## Discussion

Our previously published Poisson-multinomial model demonstrated greater accuracy in estimating intact templates from duplex (“2D”) droplet dPCR data compared with contemporaneous approaches [18,20,21,23]. Building upon this foundation, we extended the model to quadplex reactions, enabling estimation of linkage between four targets along nucleic acid templates. While triplex (“3D”) linkage analysis has been recently published, the statistical framework was not fully described [32]. Quantitation of intact viral DNA using combinatorial analyses of 5-channel multiplexed data have also been described that do not employ linkage modeling [33–35]. Here we provide full transparency on the statistical approach used to model linkage of four-channel droplet dPCR data, enabling accurate deconvolution of multi-positive droplets and we provide the R code for public use.

We tested the performance of the novel quadplex model across varied experimental conditions. In controlled plasmid experiments, the model generally produced accurate estimates of intact template probabilities for samples of either intact, bisected, or mixed template populations containing low proportions of fully fragmented material. Modeling accuracy deteriorated as the proportions of extensively fragmented templates in samples increased, particularly at higher concentrations. In these conditions, theoretically unexpected fragment types became over-represented, resulting in negative estimates of fully intact templates (p1111). These findings suggest that model performance depends on the combined effects of template fragmentation and droplet occupancy. While increased fragmentation may contribute to higher occupancy, the present study was not designed to isolate fragment-size effects at matched molar occupancy. Future studies controlling for these parameters could determine if fragment lengths may independently influence model performance. Accordingly, the empirical operating ranges reported here should be interpreted as assay- and application-specific rather than as a universally transferrable criteria for any datasets.

We also noted model performance was dependent on template identity and quadplex primer/probe panel design. For instance, two of the three linearized plasmid templates we tested returned highly accurate model estimates for intact probabilities (p1111 >90% compared to the expected 100%). In contrast, estimates for the linearized AAV2-BIIB2 plasmid were more variable resulting in an average estimate of 87% intact. Notably, the corresponding AAV2-BIIB2 quadplex panel exhibited higher low-level background signal in no template control (NTC) wells than the pLV-CMV-EGFP and AAV9-BIIB1 panels, particularly for the WPRE target. Although we did not systematically investigate whether this background contributed to the lower percentage intact estimate for the AAV2-BIIB2 linearized plasmid control, it is possible non-specific amplification, probe-associated background or low-level cross-talk may affect model performance. For accurate modeling of intact templates, we propose thorough empirical testing of model limits using vector-specific plasmid controls, and implementation of an application-specific occupancy threshold based on the empty-droplet fraction or corresponding λ value, established using representative templates and assay conditions. Several quadplex primer/probe panels should be screened to select an optimally performing set with even amplification across targets, adequate cluster separation, and minimal NTC signal.

We envision utility of the quadplex model in characterizing both viral and non-viral gene therapy vectors. For viral vectors, the model can be used for screening candidate product quality, comparing process vector designs, or process upstream and downstream modifications. As the model provides not only estimates of completely intact templates (p1111) but also all possible iterations of linkage combinations, flexible characterization of sequences depending on the analysts’ experimental question is possible. Model applications for non-viral vectors include confirmation of coding sequences or regulatory elements, and detection of aberrant sequences. Furthermore, use of the model can be extended to any nucleic acid template as multiplex RT-dPCR and linkage analyses have been successfully implemented with RNA templates [36]. Finally, this model may easily be employed to quantitate and characterize the presence of intact residual contaminating DNAs generated by the production process such as host cell or plasmid DNA.

## Materials & Methods

### Vectors

All plasmids were designed by Biogen. AAV-BIIB material was produced by Biogen using a transient transfection process.

### Primers and Probes

Custom designed primers and TaqMan QSY and QSY2 probes targeting cytomegalovirus enhancer (CMV), synapsin promoter, puromycin resistance, ampicillin resistance, WPRE, SV40pA, hBGpA, EGFP, and proprietary AAV9-BIIB1 and AAV2-BIIB2 GOIs were purchased from Thermo Fisher Scientific:

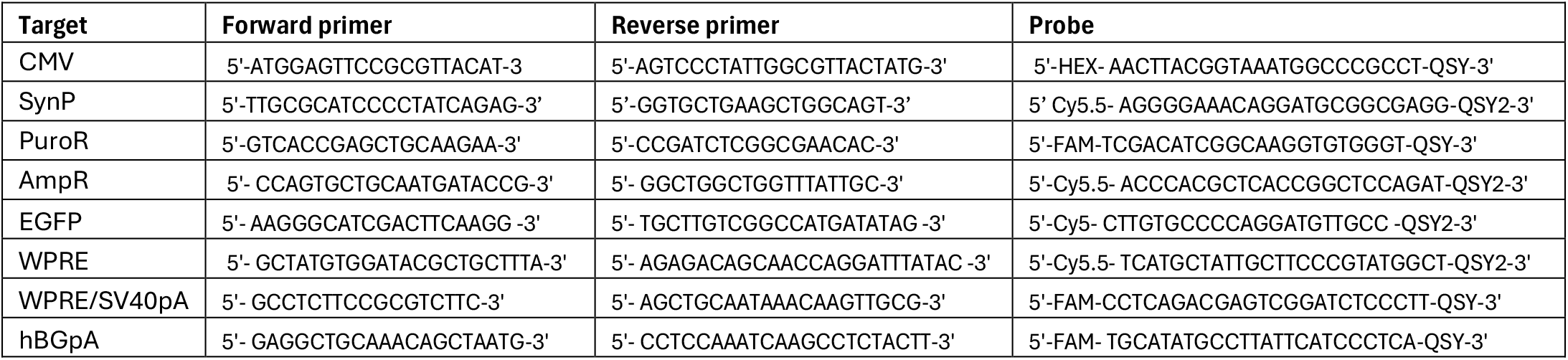

### Sample preparation

#### Digested plasmid templates

pLV-CMV-EGFP was digested with MfeI alone or varying combinations of MfeI, SnaBI, PacI and EagI. pAAV9-BIIB1 and pAAV2-BIIB2 were linearized in single digestion reactions with XhoI and MfeI respectively (New England Biolabs). The purity and theoretical DNA concentrations of digests were confirmed using a NanoDrop spectrophotometer (Thermo Fisher Scientific) and Qubit fluorometer (Invitrogen, Qubit4) and converted to copies/mL.

Digested samples were pre-diluted to 1×10^10^ copies/mL based on fluorometric measurement in TE containing a final concentration of 10 µg/ml sheared salmon sperm DNA (Thermo, AM9680). Concentrations of normalized digests were further confirmed by singleplex droplet dPCR assays. Normalized digests were then either assayed alone or mixed at varying ratios to simulate mixed populations of different sized DNA templates and run in droplet dPCR reactions.

#### Plasmid deletion constructs

Deletion constructs ΔCMV and ΔWPRE were generated by restriction enzyme double-digestion of pLV-CMV-EGFP. The CMV promoter was removed using SpeI and NheI, and the WPRE element was removed using EagI and PsiI (New England Biolabs). Digested DNA was run on a 1% agarose gel, and bands of interest were excised and purified with a gel-extraction kit (Zymo, D4001). Linear ΔCMV DNA was ligated using Quick Ligase (New England Biolabs, M2200). ΔWPRE DNA was treated with mung bean nuclease (New England Biolabs, M0250S) and then ligated with Blunt/TA Ligase Master Mix (New England Biolabs, M0367S). Two µL of each ligation reaction were transformed into chemically competent cells (New England Biolabs, C2987H) and bacterial colonies were grown over night on LB-carbenicllin plates (Teknova, L1010). Single colonies were grown overnight in LB media containing 100 µg/mL ampicillin. Plasmid DNA was isolated from bacterial cultures and purified using a Qiagen spin prep kit (cat. 27104). Successful deletion variants were confirmed by restriction digest. Samples were subsequently prepared as described under *Digested plasmid templates*.

#### Heat-treated rAAV

AAV9-BIIB1 was pre-diluted to a target concentration of 1×10^6^ copies/µL and digested with DNase I for 10 minutes at 37°C (Thermo Scientific). Viral capsids were disrupted with 1% SDS solution and incubation at low heat (10 minutes, 60°C) followed by 1 minute at 95°C). Following decapsidation, samples were incubated at 95°C for an additional 0, 5, or 15 minutes. Samples were then pre-diluted for droplet dPCR reactions to a target concentration range (5000 copies/µL).

#### UV-treated rAAV

AAV2-BIIB2 was aliquoted into four 0.5ml tubes and treated with the germicidal UV-C lamp of a biosafety cabinet for either 0, 15, 30, or 60 minutes. Viral capsids were disrupted with SDS solution and incubation at low heat (10 minutes, 60°C). Samples were then pre-diluted for droplet dPCR reactions to a target concentration range (5000 copies/µL).

### Quadplex ddPCR

Following sample preparation, samples were serially diluted in TE buffer containing 0.05% Pluronic F-68 (Gibco, 24040032), and combined with droplet dPCR master mix with or without the addition of SmaI for AAV and plasmid templates, respectively (New England Biolabs). Samples were transferred to QX ONE cartridges (Bio-Rad 12006859) and partitioned into approximately 20,000 droplets by the QX ONE droplet dPCR system.Droplets for pLV-CMV-EGFP experiments were subjected to endpoint PCR thermal cycling within the QX ONE System with thermocycler protocol: 1 cycle of 95°C × 10’; 40 cycles of 94°C × 30’’, 60°C × 1’, and 1 cycle of: 98°C × 10’). Droplets for rAAV experiments were subjected to ITR digestion with SmaI and endpoint PCR thermal cycling within the QX ONE System with thermocycler protocol: 1 cycle of 37°C × 15’; 1 cycle of 95°C × 10’; 40 cycles of 94°C × 30’’, 60°C × 1’, and 1 cycle of: 98°C × 10’). Samples were read within the Bio-Rad QX ONE droplet dPCR System and analyzed using QX ONE Software, Regulatory Edition (Bio-Rad 12012078).

### Quadplex Poisson-multinomial model

Data quality of all experiments was qualitatively assessed, and positive/negative droplets were thresholded within the QX ONE software (Bio-Rad) using amplitude plots. Identical thresholds were applied across wells within a given experiment. Following threshold assignment, droplets were automatically classified by the QX ONE software into the 16 possible 4-channel cluster states (0000-1111). Two datafiles from the QX ONE Data Table menu (Visible Data and Cluster Data) generated by the QX ONE System were exported from the instrument as separate *.csv files. Both files are required for quadplex modeling and are imported into the R-coded model. The model requires manual input of channel/target identity and inputs of ‘Accepted Droplets’ from the Visible Data export for calculation of lambda. Columns B-E of the Cluster Data export correspond to the binary readout of positive/negative (1 or 0) for amplification of each target for each fluorescent channel (Target 1-4) and Column F contains the corresponding Counts for each droplet cluster type. These droplet clusters are required for modeling fragment type probabilities referred to in the manuscript (ie amplification of all 4 targets ~ p1111). Model estimates are captured as output files with columns for all fragment type probabilities.

It is worth noting any samples with <10,000 in the raw data Accepted Droplets column are excluded by the code. In the event that a well contained no empty droplets, the maximum calculated lambda value within the plate is substituted for that well. All rAAV datasets were filtered using empirically derived criteria established through the proof of concept plasmid experiments: samples with <27.6% empty-droplet fraction and <39 copies/μl were excluded.

### Long-read NGS

rAAV DNA was extracted and annealed to form duplex templates suitable for long-read sequencing. For each sample, 2×10^13^ vg of AAV9-BIIB1 material was lysed with an SDS solution. ssDNA was annealed by rapid heating and cooling using a C1000 Touch thermocycler (Bio-Rad) to form dsDNA. Annealed templates were purified using QIAquick PCR purification kit (Qiagen). Purified DNA was measured using both a NanoDrop spectrophotometer (Thermo Fisher Scientific) and Qubit fluorometer (Invitrogen, Qubit4). 1 μg samples diluted to 133 ng/μL based on fluorometric concentration readings were sent for long-read sequencing. Sequencing was performed with a PacBio Sequel IIe following manufacturer’s guidelines for SMRTbell prep kits. A custom python-based analytical pipeline was used to quantitate %intact rates. All sequencing reads were aligned with BWA-MEM [37], low-quality alignments were removed with SAMtools [38]. Percent genome integrity was estimated using the analytical pipeline that defined “full-length reads” as >90% AAV ITR-ITR coverage, and calculated integrity as: (number full length reads)/(number of positive reads)*100.

### Protein expression assay

#### Cell transfection

Serial dilutions of AAV2-BIIB2 reference standard, control, and UV-treated samples, were plated in triplicate in 96-well plates. HEK 293T (ATCC, CRL-3216) cells were added to the serially diluted samples and assay plates were incubated for 48 hours at 37 °C with 5% CO2. All dilutions were performed in pre-warmed assay medium (DMEM+10% FBS). Three independent experimental replicates were performed on different days.

#### MSD immunoassay

Streptavidin-coated 96-well plates (MSD, L15SA) were blocked with 3% BSA for 30 minutes. Following 48-hour incubation, transfected cells were lysed with RIPA buffer containing protease and phosphatase inhibitors. Cell lysates were plated onto blocked assay plates followed by solutions of biotinylated capture antibodies and SULFO-TAG-labeled detection antibodies diluted in 1x PBS containing 1.0% BSA and1.5% Tween-20. Following 3 hours of incubation at room temperature, diluted 4x read buffer (MSD, R92TC-1) was added to each well and plates were read using MSD readers (MSD, SI 2400).

### Data analysis

All raw data, processed datasets, and R scripts used for quadplex modeling and figure generation are publicly available at: https://github.com/ddPCR/quadplex_model.

Droplet analysis was performed using Bio-Rad QuantaSoft Software, Regulatory Edition (version 1.7.4). Inter-assay replicates and experimental replicates are indicated in the figure legends. Percent recovery was calculated by comparing the calculated genome integrity to the expected genome integrity using the following equation:

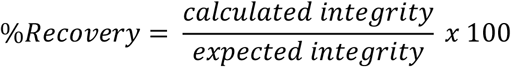

Percent relative standard deviation (RSD) was calculated as the standard deviation of the calculated integrity replicates divided by the average of the calculated integrity replicates, multiplied by 100. Assay accuracy was calculated as the grand average of the replicate average percent recovery values. Intermediate precision was calculated as the RSD of the replicate average percent recovery values. Pre-defined acceptance criteria of RSD <20% and recoveries between 80-120% were used for data analysis of all experiments.

Linear correlation of average modeled probability of intact templates (p1111) and maximal protein expression (Emax) calculated in Prism was assessed using Pearson’s correlation coefficient in R. Simple linear regression was used for visualization of the relationship. As measurements represent averages of replicates, a linear mixed-effects model including replicate as a random effect was evaluated. Results were consistent with simple linear regression, statistical outputs are provided in accompanying R scripts.

## Acknowledgments

We thank our colleagues in Biogen Analytical Development for their scientific discussions and technical support. We are grateful to Svetlana Bergelson for her analytical review and resource management. We sincerely thank Thomas Carlile and Wei Li for their NGS support. Their efforts were instrumental in the timely generation and interpretation of sequencing data during manuscript revision.

