## Supplemental figures for "A Poisson-multinomial model for quantifying gene therapy vector integrity using quadplex digital PCR"

### Supplemental information

**S1 Fig. Plasmid 1D amplitude plots.** 1D amplitude plots for each fluorescent channel with optimal primer/probe set corresponding to Fig 1 (CMV:HEX, EGFP:Cy5, PuroR:FAM, AmpR:Cy5.5,well A2, expected 2500 copies/µL).

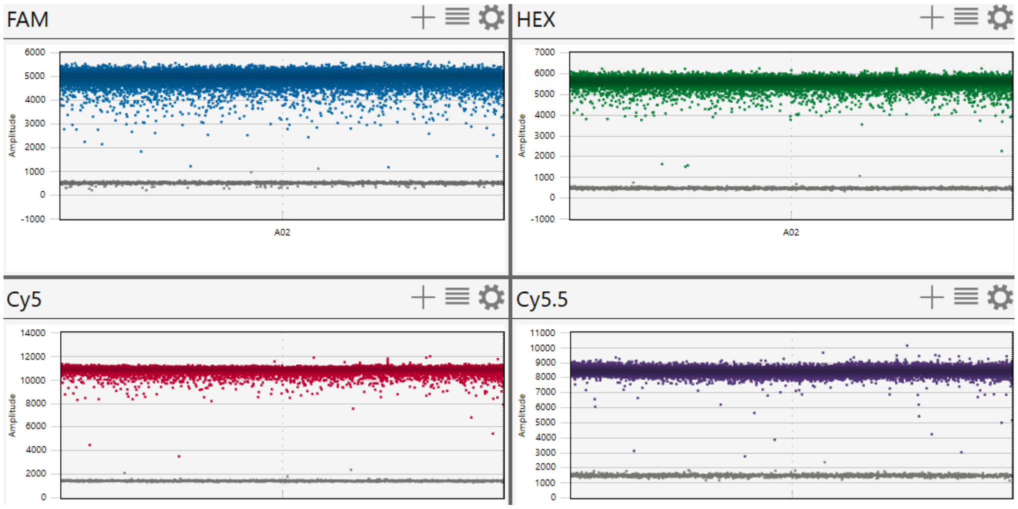


**S2 Fig. Plasmid 2D amplitude plots.** 2D amplitude plots for all pairs of fluorescent channels with optimal primer/probe set corresponding to Fig 1 (CMV:HEX, EGFP:Cy5, PuroR:FAM, AmpR:Cy5.5, well A2, expected 2500 copies/µL).

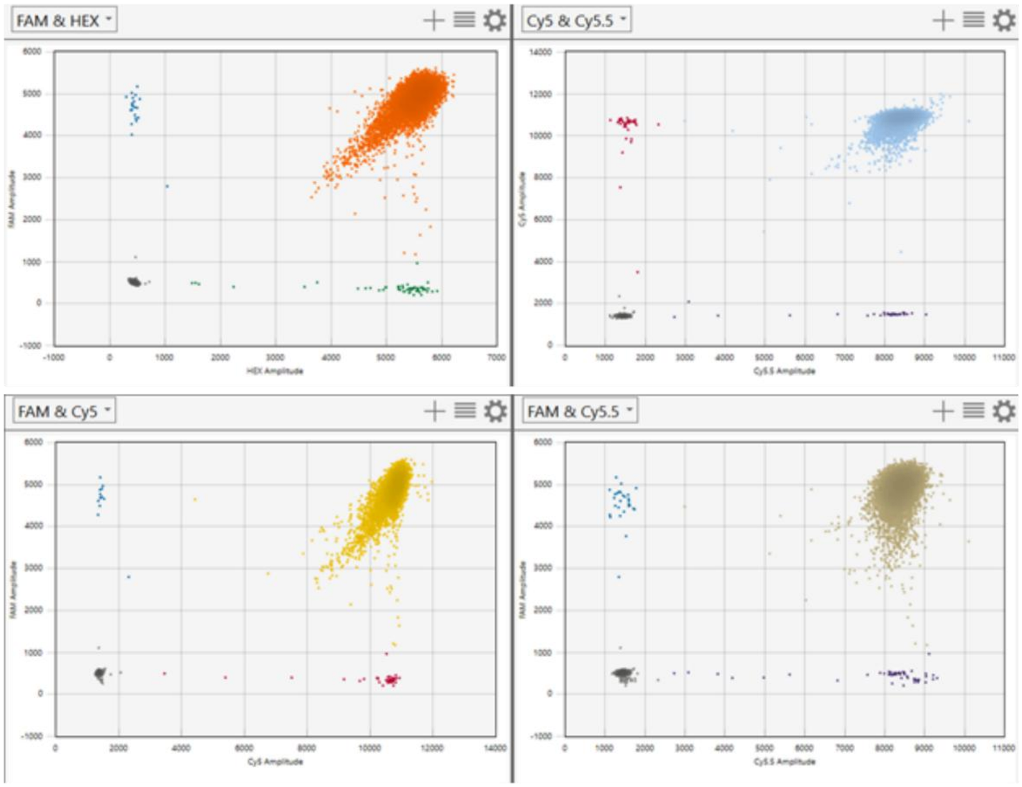

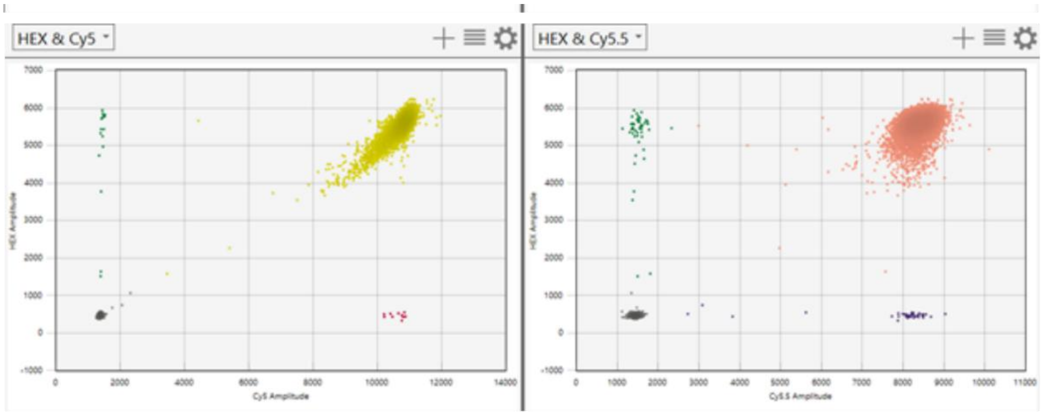


**S3 Fig. Plasmid Negative Control droplet plots.** 1D amplitude plots for each fluorescent channel of negative template control corresponding to Fig 1 (CMV:HEX, EGFP:Cy5, PuroR:FAM, AmpR:Cy5.5, all <2copies/μL).

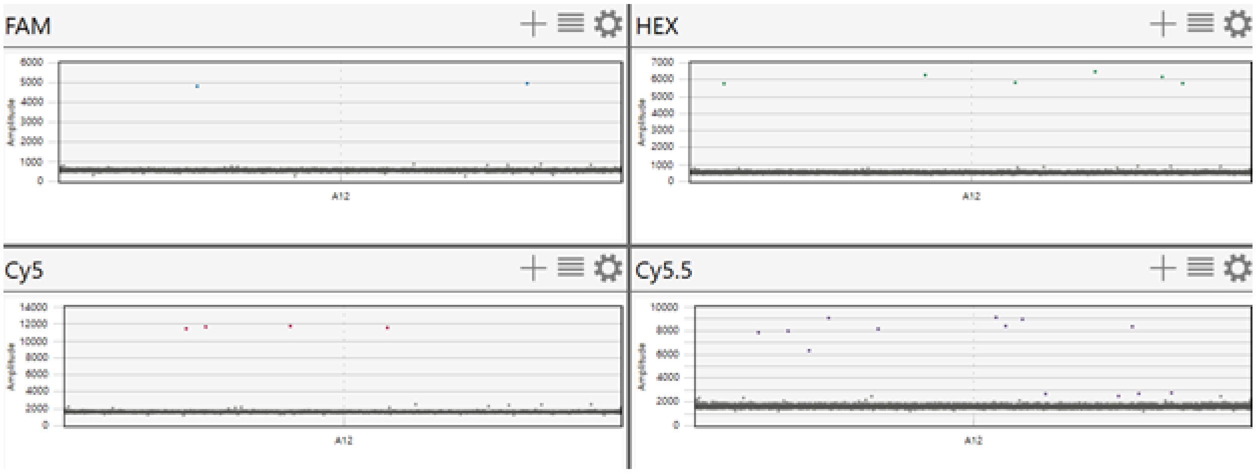


**S4 Fig. 1D amplitude plots of alternative (non-optimal) quadplex primer/probe panel.** 1D amplitude plots for each fluorescent channel with alternative primer/probe set: CMV-FAM, CAG-VIC, EGFP-Cy5, WPRE-Cy5.5. This alternative panel demonstrated reduced droplet separation and droplet cluster distortion consistent with inter-channel signal interference. Sample pLV-CMV-EGFP MfeI digest, well D1, expected 1700 copies/μL.

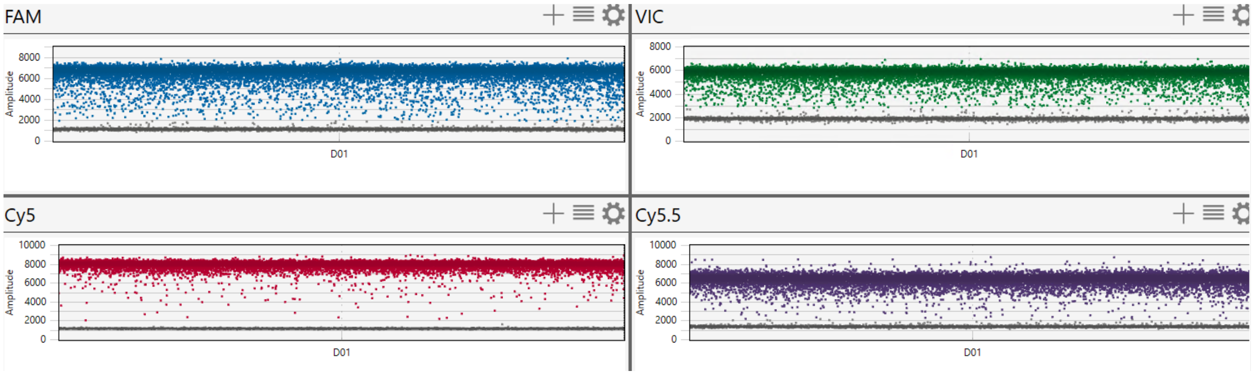

**S5 Fig. 2D amplitude plots of alternative (non-optimal) quadplex primer/probe panel.** 2D amplitude plots for each fluorescent channel with alternative primer/probe set: CMV-FAM, CAG-VIC, EGFP-Cy5, WPRE-Cy5.5. This alternative panel demonstrated reduced droplet separation and droplet cluster distortion consistent with inter-channel signal interference. Sample pLV-CMV-EGFP MfeI digest, well D1, expected 1700 copies/μL.

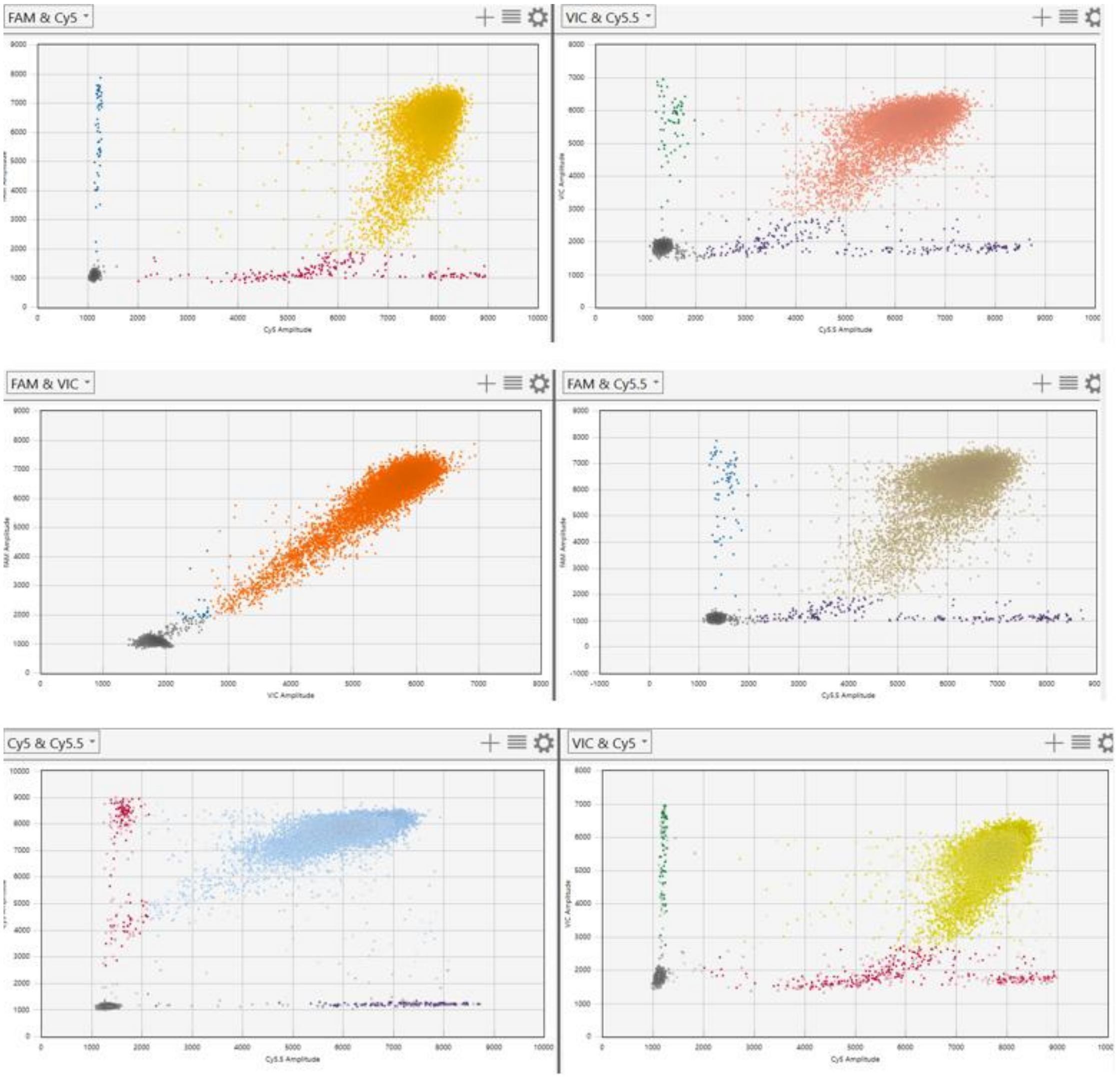


**S6 Fig. NTC plots of alternative (non-optimal) quadplex primer/probe panel.** 1D amplitude plots for each fluorescent channel of no template control with alternative primer/probe set: CMV-FAM, CAG-VIC, EGFP-Cy5, WPRE-Cy5.5, all <3 copies/μL).

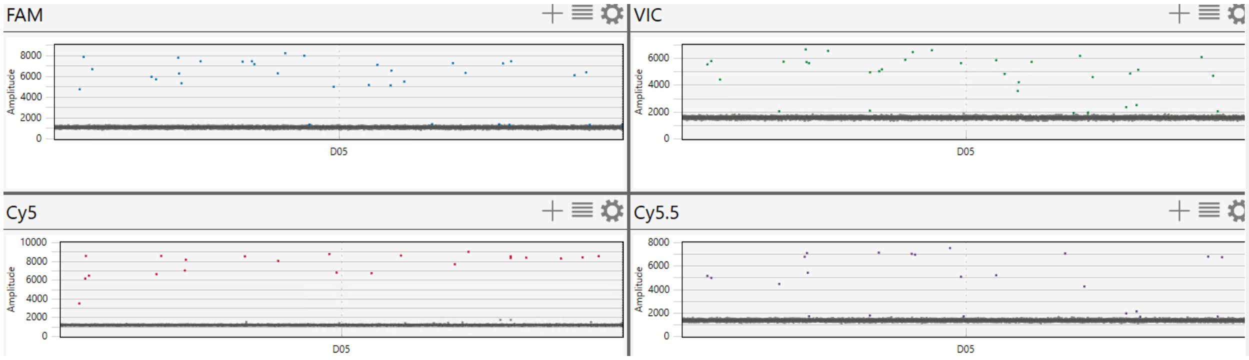

**S7 Fig. Plasmid sample preparation.** pLV-CMV-EGFP digests were run on a 1% agarose gel and visualized with SYBR Safe (Invitrogen) on an Odessey M imaging system (Licor). From left to right: Lane 1: NEB 1KB ladder (catalog N0552S); Lane 2: MfeI; Lane 3: MfeI and PacI; Lane 4: MfeI and EagI; Lane 5: MfeI, PacI, and EagI (each corresponds to Fig. 3A,C,D,G respectively).

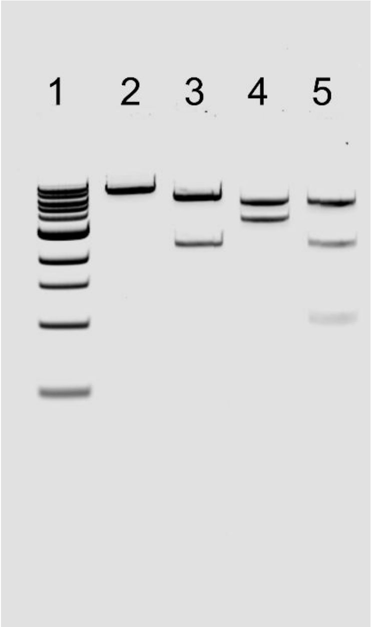


**S8 Fig. Plasmid sample preparation (continued).** pLV-CMV-EGFP digests were run on a 1% agarose gel and visualized with SYBR Safe (Invitrogen) on an Odessey M imaging system (Licor). From left to right: Lane 1: NEB 1KB ladder (catalog N0552S); Lane 2: MfeI and SnaBI; Lane 3: MfeI, SnaBI and PacI; Lane 4: MfeI, SnaBI and EagI; Lane 5: MfeI, SnaBI, PacI, and EagI (each corresponds to Fig 3B, E, F, H respectively).

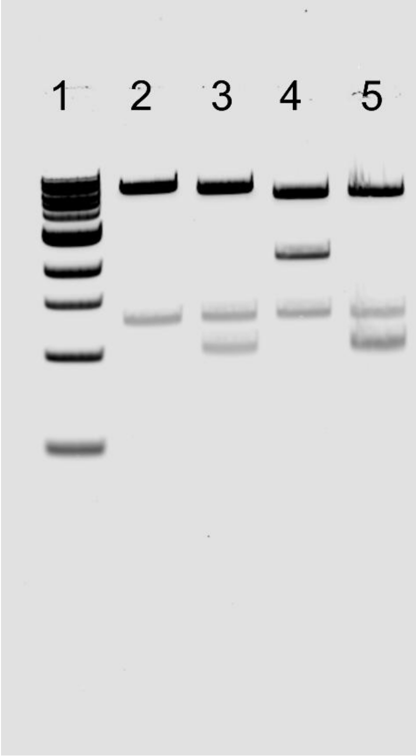

**S9 Fig. Model results for pLV-CMV-EGFP MfeI digests with alternative (non-optimal) quadplex primer/probe panel.** Average model estimates of fragment type probabilities. Mean intact probability = 93.2% tested at 850-1700 copies/µL.

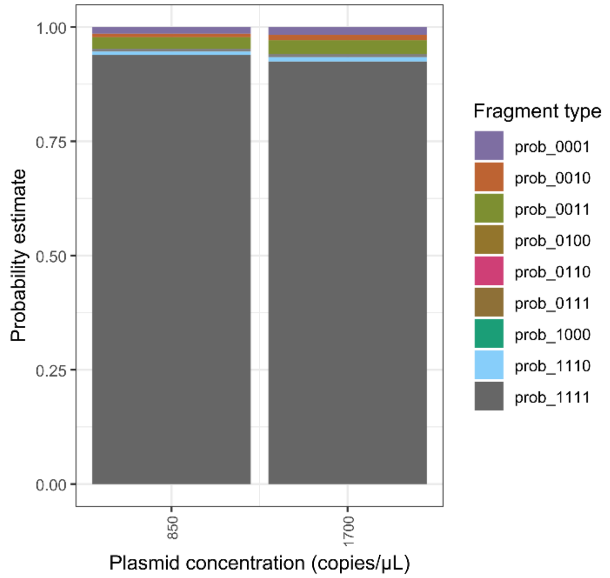


**S10 Fig. Percent recovery of measured vs expected probabilities for each expected fragment type presented in Table 2.** Points represent individual dilution values per sample and are color coded by dilution concentration. Bars represent the average of all dilutions per sample. Error bars signify standard deviation.

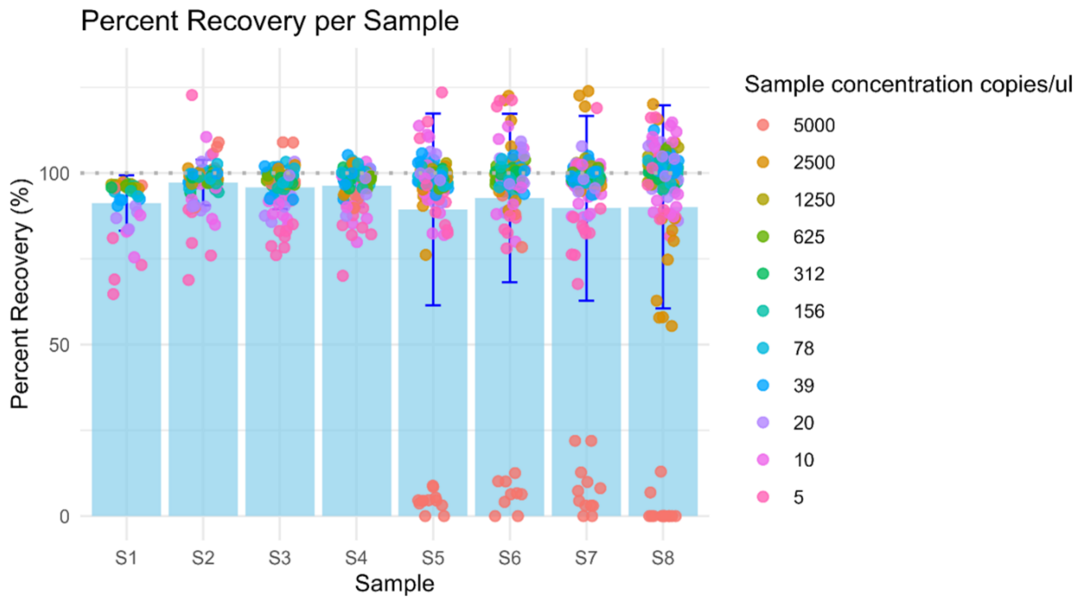

**S11 Fig. Percent recovery of measured vs expected probabilities for each expected fragment types presented in Tables 3 and 4.** Samples with expected fragment probabilities of zero cannot be used to calculate percent recovery and were excluded from plot. Points represent individual dilution values per sample and are color coded by dilution concentration. Bars represent the average of all dilutions per sample. Error bars signify standard deviation.

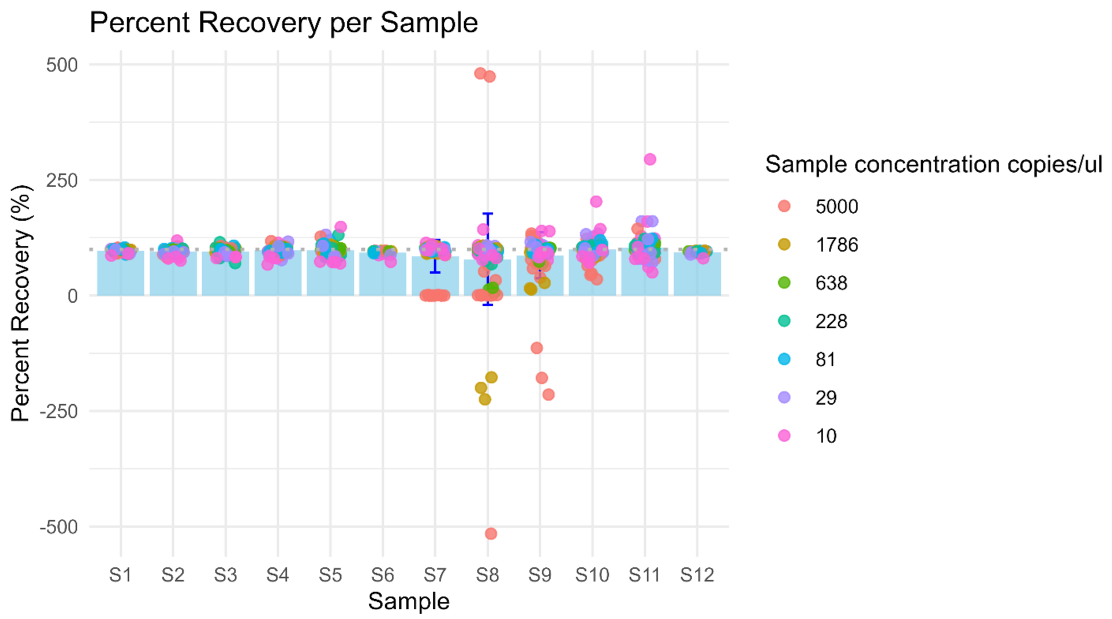


**S1 Table. Sample preparation of CMV deletion variant mixtures.** Samples prepared as mixtures of pLV-CMV-EGFP MfeI digests (100% intact) and ΔCMV deletion variant linearized with MfeI and their expected target amplification probabilities.

| Sample Number | pLV-CMV-EGFP MfeI digest (%) | ΔCMV (%) | Expected p1111 |
|---|---|---|---|
| Sample 1 | 100 | 0 | 100 |
| Sample 2 | 75 | 25 | 75 |
| Sample 3 | 50 | 50 | 50 |
| Sample 4 | 25 | 75 | 25 |
| Sample 5 | 0 | 100 | 0 |

**S2 Table. Sample preparation of WPRE deletion variant mixtures.** Samples prepared as mixtures of pLV-CMV-EGFP MfeI digests (100% intact) and ΔWPRE deletion variant linearized with MfeI and their expected target amplification probabilities.

| Sample Number | pLV-CMV-EGFP MfeI digest (%) | ΔWPRE (%) | Expected p1111 |
|---|---|---|---|
| Sample 6 | 100 | 0 | 100 |
| Sample 7 | 75 | 25 | 75 |
| Sample 8 | 50 | 50 | 50 |
| Sample 9 | 25 | 75 | 25 |
| Sample 10 | 0 | 100 | 0 |

**S12 Fig. Model results for mixed deletion variant samples.** Panels A-E correspond to ΔCMV samples 1-5 left to right. Panels G-K correspond to ΔWPRE samples 6-10 left to right, n=2.

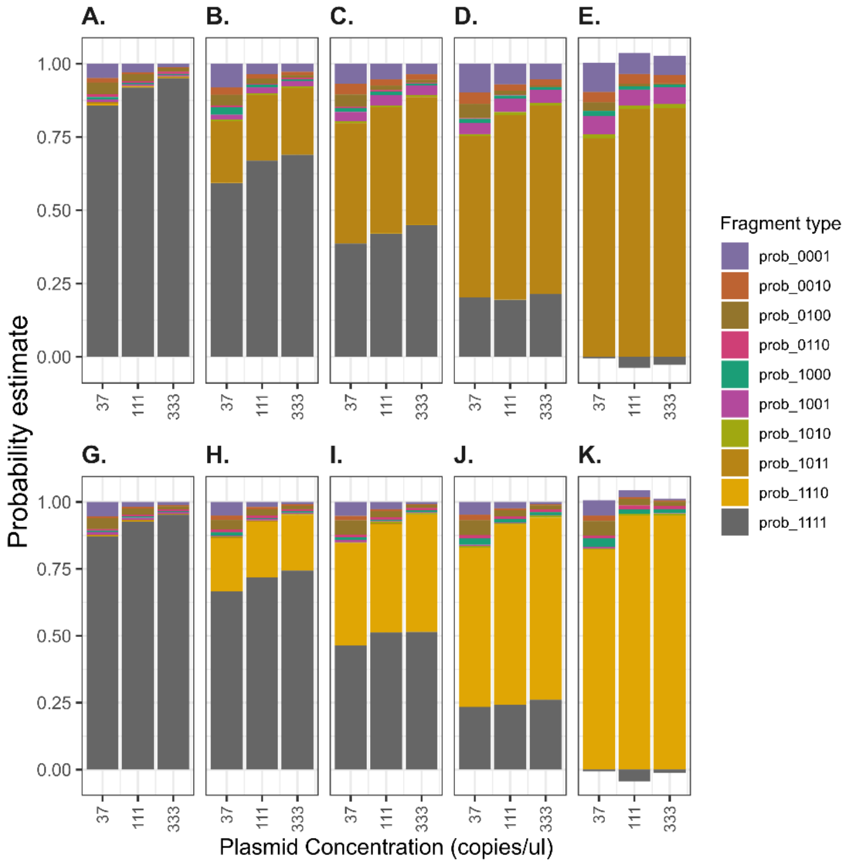


**S3 Table. Summary results of model estimates for deletion variant mixtures.** Average model estimates of percent intact and fragmented genomes corresponding to S12 Fig, n=2.

| | Modeled %intact genomes | | |
|---|---|---|---|
| Sample | p1111 | p1011 | p1110 |
| S1 | 90.9% | 0.4% | 0.4% |
| S2 | 65.0% | 22.0% | 0.2% |
| S3 | 41.9% | 42.4% | 0.1% |
| S4 | 20.4% | 60.8% | 0.0% |
| S5 | -2.3% | 81.3% | 0.0% |
| S6 | 91.7% | 0.3% | 0.2% |
| S7 | 70.9% | 0.3% | 20.5% |
| S8 | 49.3% | 0.2% | 40.3% |
| S9 | 24.6% | 0.0% | 65.0% |
| S10 | -2.1% | 0.0% | 90.7% |

**S13 Fig. AAV9-BIIB1 vector and primer/probe diagram**. (P1 = FAM, P2 = VIC, P3 = Cy5, P4 = CY5.5).

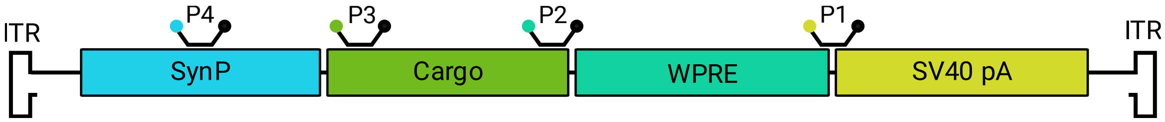


**S14 Fig. AAV2-BIIB2 vector and primer/probe diagram**. (P1 = FAM, P2 = HEX, P3 = Cy5, P4 = CY5.5).

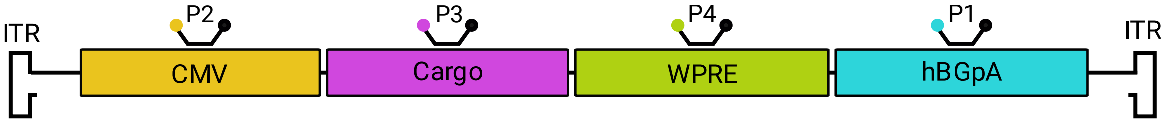


**S15 Fig. AAV9-BIIB1 1D amplitude plots.** 1D amplitude plots for each fluorescent channel with optimal primer/probe set corresponding to S13 Fig (SynP:Cy5.5, 5'GOI:Cy5, 3'GOI:VIC, WPRE/SV40:FAM), well C6, expected 800 copies/μL.

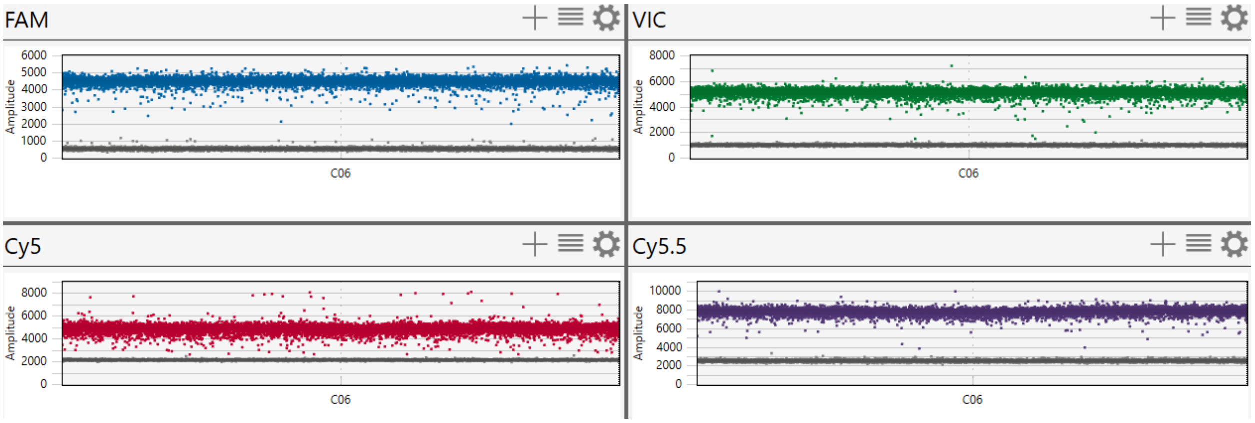

**S16 Fig. AAV9-BIIB1 2D amplitude plots.** 2D amplitude plots for each fluorescent channel with optimal primer/probe set corresponding to S13 Fig (SynP:Cy5.5, 5’GOI:Cy5, 3’GOI:VIC, WPRE/SV40:FAM), well C6, expected 800 copies/μL.

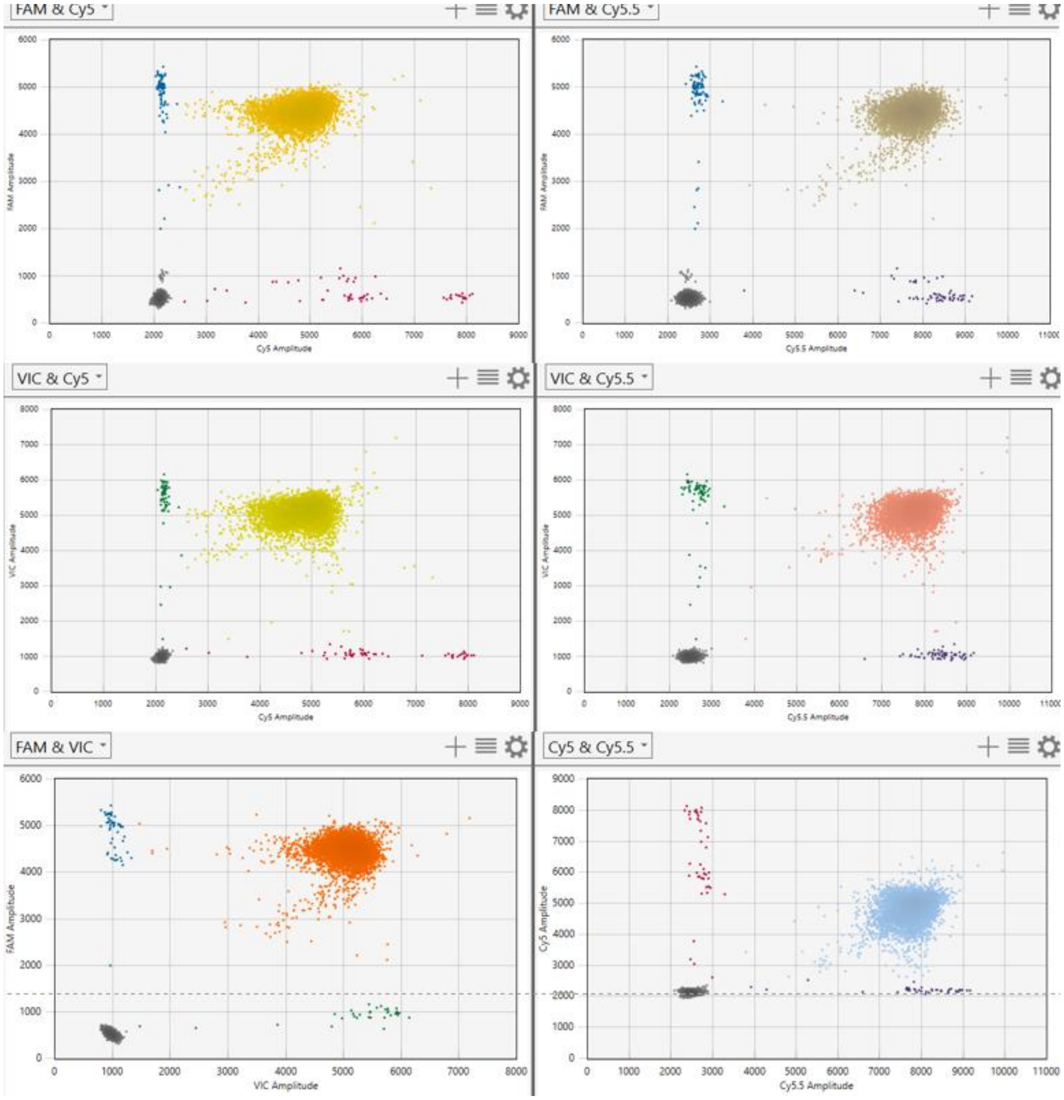


**S17 Fig. AAV9-BIIB1 NTC amplitude plots.** 1D amplitude plots for each fluorescent channel of no template control with AAV9-BIIB1 quadplex panel corresponding to S13 Fig (SynP:Cy5.5, 5’GOI:Cy5, 3’GOI:VIC, WPRE/SV40:FAM) (all <2copies/μL).

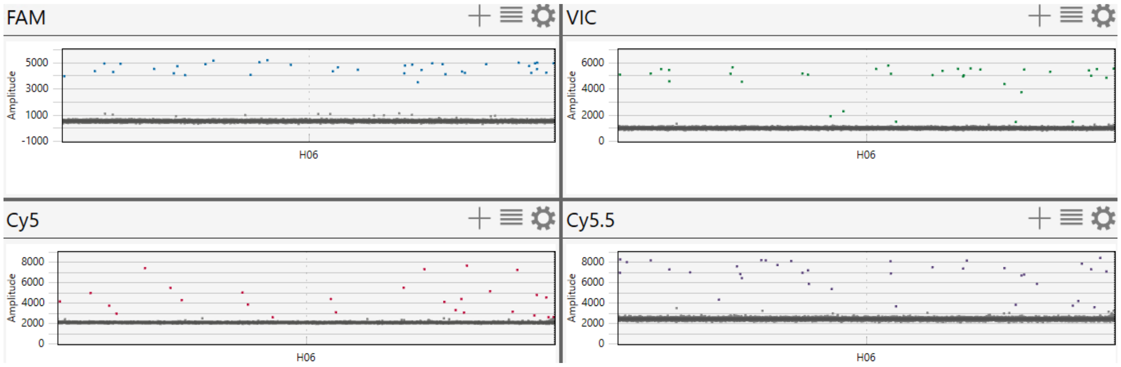

**S18 Fig. AAV2-BIIB2 1D amplitude plots.** 1D amplitude plots for each fluorescent channel with optimal primer/probe set corresponding to S14 Fig (CMV:HEX, GOI:Cy5, WPRE:Cy5.5, hBGPA:FAM), well C1, expected 556 copies/μL.

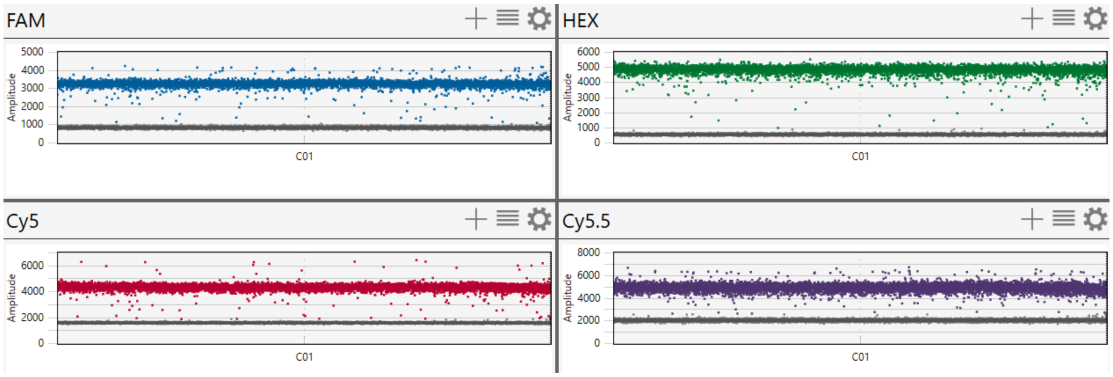


**S19 Fig. AAV2-BIIB2 2D amplitude plots.** 2D amplitude plots for each fluorescent channel with optimal primer/probe set corresponding to S14 Fig (CMV:HEX, GOI:Cy5, WPRE:Cy5.5, hBGPA:FAM), well C1, expected 556 copies/μL.

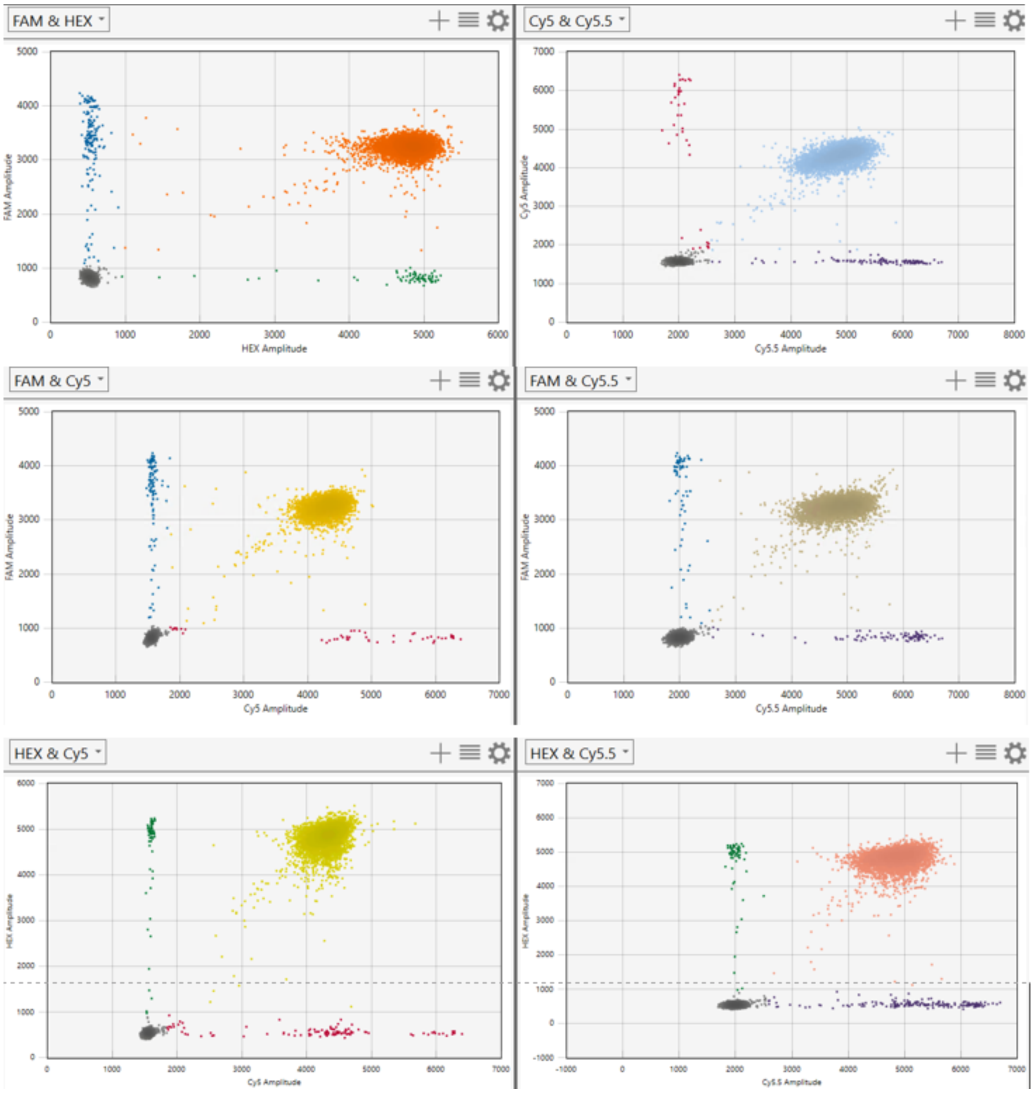

**S20 Fig. AAV2-BIIB2 NTC amplitude plots.** 1D amplitude plots for each fluorescent channel of no template control with AAV2-BIIB2 quadplex panel corresponding to S14 Fig (CMV:HEX, GOI:Cy5, WPRE:Cy5.5, hBGPA:FAM), (all <20copies/µL).

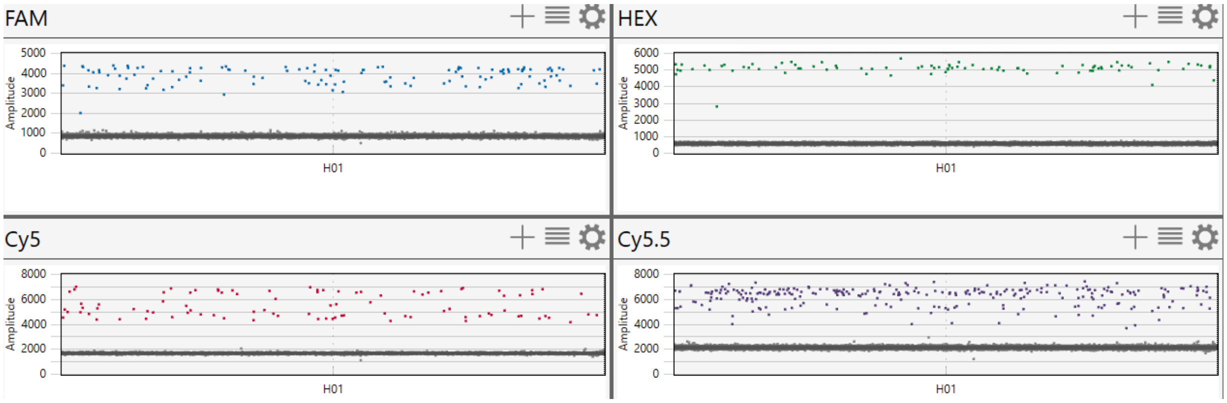


**S4 Table. Comparison of long-read NGS and quadplex model integrity estimates.** Extracted ssAAV9-BIIB1 genomes were annealed and purified to generate duplex templates suitable for sequencing. The same purified duplex samples were analyzed by long-read NGS and by quadplex droplet dPCR followed by quadplex model analysis. Resulting estimates of percentage of intact genomes are shown for method comparison.

| Material | Condition | intact rate NGS (%) n=1 | intact rate ddPCR quadplex model of NGS samples (% n=1) |
|---|---|---|---|
| AAV9-BIIB1 | Control | 87 | 68 |
| | 5 min 95°C | 78 | 59 |
| | 15 min 95°C | 38 | 22 |

**S21 Fig. Long-read NGS samples visualized with gel electrophoresis.** Extracted and annealed dsDNA templates for control and stressed AAV9-BIIB1 were run on a 1% agarose gel and visualized with SYBR Safe (Invitrogen) on an Odessey M imaging system (Licor). From left to right: Lane 1: NEB 1KB ladder (catalog N0552S); Lane 2 AAV9-BIIB1 Control; Lane 3: AAV9-BIIB1 5 minutes 95°C; Lane 4: AAV9-BIIB1 15 minutes 95°C.

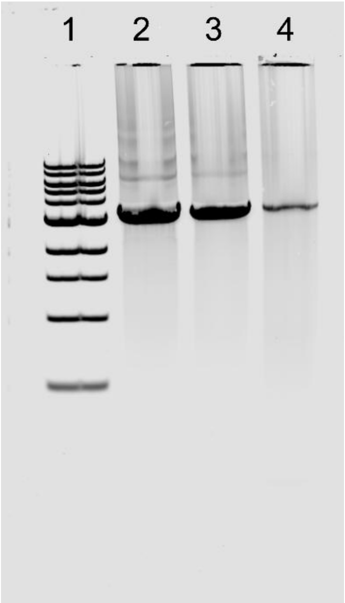
